# Asymmetric migration shapes genetic structure of the invasive avian vampire fly (*Philornis downsi*) across the Galápagos Islands

**DOI:** 10.64898/2026.07.03.735711

**Authors:** Abbie C. Hay, Sonia Kleindorfer, Lauren K. Common, Sally Potter, Jennifer A.H. Koop, George E. Heimpel, Sarah A. Knutie, Birgit Fessl, Lorraine Pérez-Beauchamp, Rachael Y. Dudaniec

## Abstract

Biological invasions on islands provide a natural framework to study how dispersal and connectivity influence evolutionary and ecological processes. The avian nest parasitic fly, *Philornis downsi –* first recorded in Darwin’s finch nests in 1997 – causes high mortality in endemic land birds, yet its inter-island and sex-specific patterns of dispersal and genetic structure remain poorly understood. We use low-coverage whole genome sequencing to investigate genome-wide patterns of genetic diversity, directional migration and effective population size in *P. downsi* across five major Galápagos Islands and its native range in mainland Ecuador. We find evidence for a genetic bottleneck in the Galápagos, isolation by distance, and evidence that the island closest to the Ecuadorian mainland, San Cristóbal, is genetically divergent from the other four islands sampled, despite retaining the highest genetic diversity. No evidence was found for sex-biased dispersal; however, sex-biased genetic structure was detected using only markers from inferred autosomal scaffolds. We found asymmetric gene flow with higher migration rates from San Cristóbal westward to the other islands, matching the direction of both southeast trade winds and major cargo shipping routes. Our results suggest both natural and human-mediated colonisation of *P. downsi* from the mainland through San Cristóbal to the other islands, followed by high inter-island dispersal among closely situated sink islands. Our findings are critical for prioritising islands for control strategies that will reduce *P. downsi* impacts on vulnerable endemic birds and underscore the value of understanding directional migration patterns for managing invasive species in metapopulations.

## Introduction

Biological invasions are a major threat to biodiversity and ecosystem health worldwide (Bellard et al. 2016; Pyšek et al. 2020; Christian 2023). Insects are particularly important invaders due to their significant ecological and economic impacts (Bradshaw et al. 2016; Pimentel et al. 2005) and strong dispersal abilities (Ochocki and Miller, 2017; Renault et al. 2018) that facilitate rapid range expansion (Seebens et al. 2017). While many insects can disperse long distances, their dispersal strategies remain poorly understood (Asplen, 2018). Insects may enter new habitats through natural vectors such as winds and passive drift on currents, or anthropogenic pathways, including maritime shipping and trade networks (Hulme et al. 2008; Wilson et al. 2009; Gippet et al. 2019). Identifying dispersal routes is logistically challenging, particularly for small-bodied insects that evade early detection (Osborne et al. 2002; Lushai and Loxdale, 2004; Liebhold et al. 2016). In such cases, invasion pathways can be reconstructed by integrating abiotic processes, geographic barriers, anthropogenic trade networks, and population genetic patterns (Estoup and Guillemaud, 2010; Cristescu 2016; Hudson et al. 2022).

Genetic approaches can track invasive species by uncovering landscape-scale patterns of neutral genetic variation shaped by both historical colonization (Dlugosch and Parker, 2008) and ongoing movement (Broquet and Petit, 2009). Demographic events – including founding events and bottlenecks – leave lasting genetic signatures such as reduced genetic diversity and increased genetic differentiation in isolated populations, potentially impacting fitness (Taylor and Keller, 2007; Excoffier et al. 2009; Sjölund et al. 2019). Many invasive species thrive despite severe bottlenecks, i.e. the ‘genetic paradox of invasion’ (Estoup et al. 2016), often through local adaptation (e.g. Kardum Hjort et al. 2024). Conversely, genetic diversity can remain high when repeated introductions from divergent lineages are admixed (Kolbe et al. 2004; Sherpa et al. 2019; Kołodziejczyk et al. 2025). Understanding these genetic processes is vital for interpreting ongoing colonization, population expansion or contraction, and forecasting control impacts.

Beyond genetic diversity, assessing the direction and magnitude of gene flow can identify processes driving invasion pathways and reveal how genetic variation is maintained. Asymmetries in gene flow can point to founding source populations that contribute disproportionately to invasion spread and increase genetic diversity of recipient sinks (Kawecki and Holt, 2002; Furrer and Pasinelli, 2016; Garnas et al. 2016). Directional gene flow has revealed invasion routes across diverse taxa, including amphibians (Everts et al. 2025), mammals (Li et al. 2024), plants (Hirata et al. 2023), and insects (Parvizi et al. 2023; Rane et al. 2023). At the individual level, sex-specific dispersal can create asymmetries where males and females differentially contribute to gene flow, generating sex-biased genetic patterns (Prugnolle and De Meeus, 2002; Blyton et al. 2015; Dudaniec et al. 2022). In such systems, differences in movement patterns between the more dispersive and the more philopatric sex can produce allele frequency variation, resulting in pronounced sex-specific signatures in bi-parentally inherited genetic markers (Goudet et al. 2002). Understanding dispersal asymmetries is critical to predicting invasion trajectories and targeting management actions.

Island archipelagos offer informative contexts to study the interplay of genetic dispersal and the impacts of invaders, particularly invasive parasites and pathogens. Invasive species are a leading threat to island biodiversity (Clavero and García-Berthou, 2005; Szabo et al. 2012; Spatz et al. 2023) where impacts on native biodiversity can be heightened due to geographic isolation (Whittaker and Fernández-Palacios, 2007), high species endemism (Kier et al. 2009), and lower genetic diversity compared to mainland populations (Frankham 1997). Invasive parasites and pathogens can be particularly damaging on islands (Wikelski et al. 2004; Causton et al. 2006; Harvey-Samuel et al. 2021), as isolated native species often lack evolved defences (Simberloff 1995; Russell et al. 2017) and exhibit altered immune systems compared to continental relatives (Matson 2006; Lobato et al. 2017; Barthe et al. 2022), which may respond differently to novel invaders. Archipelagos, with their discrete geographic boundaries and replicated metapopulation structure, are natural laboratories for investigating how dispersal and colonisation history shape genetic connectivity and population divergence (Cronk 1997; Gillespie and Roderick, 2002). These systems provide valuable insights into the invasive pathogen dynamics that threaten isolated endemic species.

The avian vampire fly, *Philornis downsi* (Diptera: Muscidae) Dodge and Aitken, 1968, is an invasive avian ectoparasite likely introduced to the Galápagos archipelago from its native range on mainland South America prior to 1964 via human-assisted dispersal (Fessl et al. 2001, 2018). First identified in the nests of endemic Darwin’s finches in 1997 (Fessl et al. 2001), free-living larvae feed on the blood and tissue of developing chicks, and occasionally nesting mothers, across a broad range of land bird species (Fessl et al. 2006; Huber et al. 2010; Kofler et al. 2025a). *Philornis downsi* is considered the greatest threat to Galápagos land birds (Causton et al. 2006; McNew and Clayton, 2018), causing up to 100% nestling mortality (O’Connor et al. 2014; Kleindorfer and Dudaniec, 2016). Since its introduction, *P. downsi* has been recorded on 15 of 17 Galápagos islands studied (Wiedenfeld et al. 2007; Fessl et al. 2018) and are known to attack at least 11 species of Darwin’s finches and other endemic bird species (Fessl et al. 2018). Adults exhibit sex-biased asymmetries in behaviour (Kleindorfer et al. 2016; Moreno-Mejía et al. 2024), dispersal (Common et al. 2022), and morphology (Common et al. 2020); however, whether sex-biased inter-island dispersal or genetic patterns exist remains unknown. Genetic studies show *P. downsi* is highly dispersive with ongoing inter-island gene flow (Dudaniec et al. 2008; Koop et al. 2021; Common et al. 2023), with emerging evidence for adaptive genetic signatures that may facilitate invasion success (Basnet et al. 2025). Mainland Ecuador is considered the most likely source location, however dispersal routes and vectors of movement that sustain ongoing inter-island connectivity remain unclear (Bulgarella et al. 2015; Fessl et al. 2018; Koop et al. 2021; Basnet et al. 2025).

Here, we explore genetic structure and dispersal in *P. downsi* across the Galápagos Islands to inform control strategies aimed at conserving endemic landbirds threatened by this ectoparasite. With low-coverage whole genome sequencing of *P. downsi* sampled across five of the Galápagos Islands we aim to; 1) characterise neutral genetic diversity, genetic structure and effective population size, with comparison to Ecuadorian mainland sites, and 2) estimate inter-island genetic migration and test for sex-biased patterns. This is the first study using whole genomic data to examine asymmetrical migration, sex-specific gene flow and effective population size in *P. downsi* across major islands of the Galápagos. Our findings will assist in prioritising locations for potential biological control or other management strategies for *P. downsi* across islands and provide insight into the role of gene flow and migration in biological island invasions.

## Materials and Methods

### Study system

*Philornis downsi* samples were collected from five major islands across the Galápagos archipelago that varied in area and human population size: Isabela (IS; 4588 km²; pop. 2,344), Santa Cruz (SZ; 986 km²; pop. 15,701) San Cristóbal (SL; 558 km²; pop. 7,088) Floreana (FL; 173 km²; pop. 111), with one island, Santiago, lacking any human settlement (SO; 585 km²; pop. 0) (Snell et al. 1996; INEC 2016) (Figure 1). Samples were collected from arid lowland habitats (0–200 m asl) on all islands, and from either humid highland zones (FL, SO, IS; 350–1000 m asl) or, where highlands were inaccessible, from adjacent agricultural areas (SZ, SL; 200–400 m asl).

**Figure 1:**
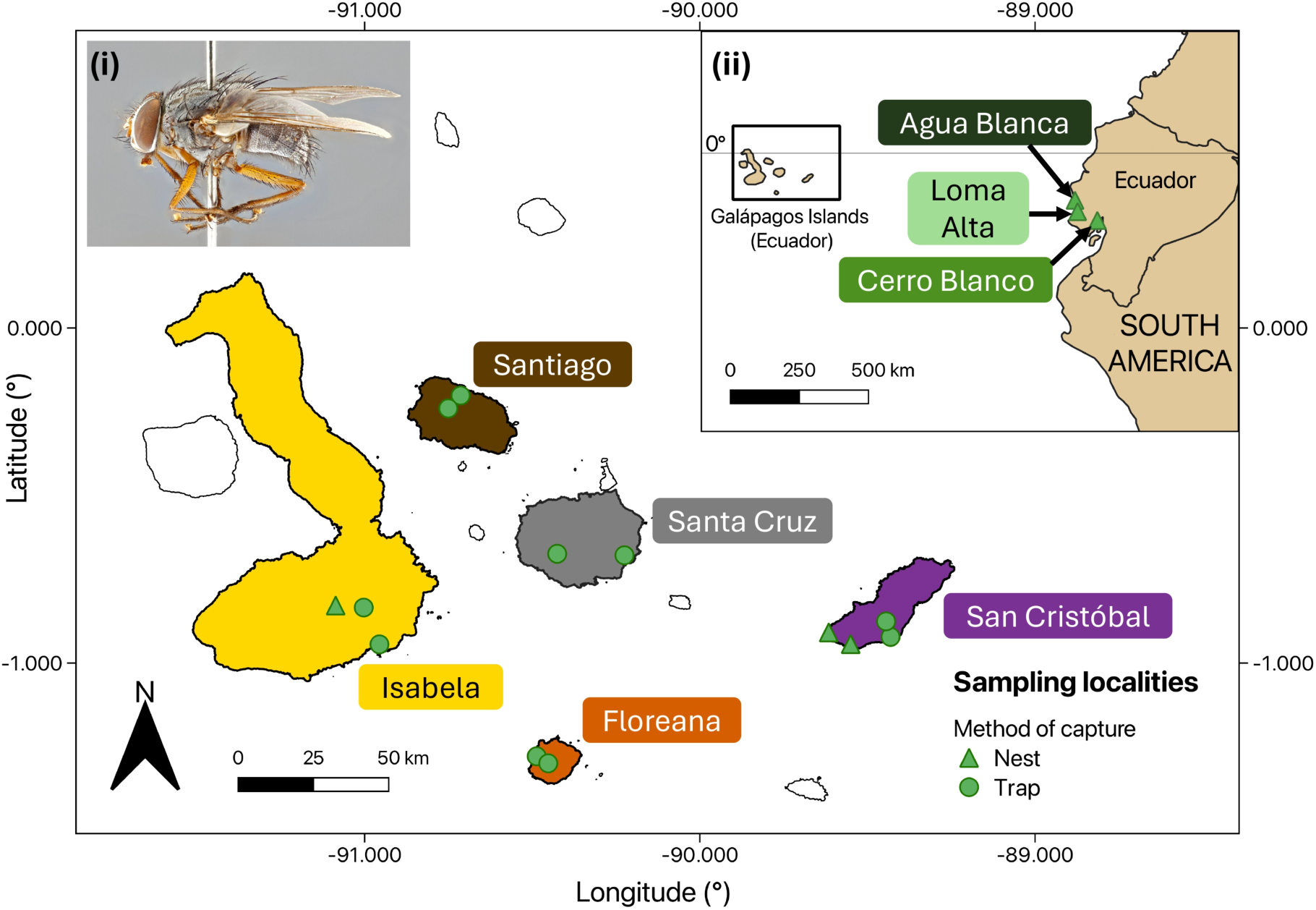
Sampling localities of *P. downsi* across five Galápagos Islands and the Ecuadorian mainland. Symbols of sampling localities correspond to the method of specimen capture (triangle = collected from terminated avian nests, circle = collected via traps). Inset (i) shows an adult male of the study species *P. downsi* (photo by B. Sinclair). Inset (ii) depicts three sampling localities on the Ecuadorian mainland, in relation to the Galápagos archipelago.

It is reported that *P. downsi* existed natively on mainland Ecuador before invading the Galápagos (Bulgarella et al. 2015), but the exact South American source population is unknown; thus, we included mainland locations as a reference native population for comparison. Samples originated from three locations on the Ecuadorian mainland – the Cerro Blanco Protected Forest, on the western edge of Guayaquil, Guayas Province (CB; 25 m asl); the Agua Blanca community within Machalilla National Park, Manabí Province (AB; 200 m asl); and the Loma Alta Ecological Reserve adjacent to the village of El Suspiro, Santa Elena province (LA; 70 m asl) (Figure 1).

### Field sampling

All research for this study complies with applicable laws on sampling from natural populations and animal experimentation. Adult flies were collected across the Galápagos Islands during the 2024 land bird breeding season (January-April) via McPhail traps (BioQuip Products, California, USA). Fly pupae were also collected from terminated nests, and pupae were reared to the adult stage where possible using protocols similar to those described in Knutie et al. (2024) and Common et al. (2025), with non-emerged individuals preserved as pupae. Samples were preserved in 80% ethanol and stored in a standard -18 °C freezer within 3 hours of collection. The species and sex of adult Galápagos specimens were identified in the field and verified via high-resolution morphological imaging for a companion study (see Figure S1). On mainland Ecuador, samples were collected as pupae from terminated nests (2015–2024) and reared to adulthood prior to species and sex identification using microscopy, as described in Bulgarella et al. (2015). See Text S1 for detailed field sampling approaches across locations. We collected 39 nest-reared adult flies from the mainland and 235 individuals from the Galápagos (138 trapped, 97 nest-derived). We sampled a varying proportion of the parasites present within each nest (3.7–100% of total nest infestation); see Text S1 for details.

### DNA extraction and low-coverage whole-genome sequencing

DNA was extracted from adult and pupal tissues using a Qiagen DNeasy Blood and Tissue kit with a modified protocol (see Text S2). Libraries were prepared by the Australian Genome Research Facility (AGRF; Melbourne, Australia) and sequenced (paired-end, 2×150 bp, low coverage), with 65–96 samples per lane across four lanes of an Illumina NovaSeq X Plus 10B (300 cycle).

### Bioinformatics and SNP calling

Raw *P. downsi* sequences were bioinformatically processed as two datasets: one combining Galápagos and mainland Ecuador samples, and one exclusively containing Galápagos samples. Quality control was performed using FASTQC v0.12.1 (Andrews 2010). Reads were adapter clipped and quality-trimmed with Trimmomatic v0.39 (Bolger et al. 2014) and Fastp v0.24.0 (Chen et al. 2018), excluding reads with Phred <20 or length <30 bp. Reads were mapped to a female *P. downsi* reference genome (GCA_019455685.1; Romine et al. 2022) using NextGenMap (Sedlazeck et al. 2013), retaining mapping quality >20 and excluding individuals with mapping efficacy <50%. Deduplication was performed in Picard v2.18.29 (Broad Institute 2019), overlapping reads were clipped using BamUtil v1.0.15 (Jun et al. 2015), and read depth was estimated using SAMtools depth v1.19.2 (Li et al. 2009).

Identification of single nucleotide polymorphisms (SNPs), genotype likelihood estimation, and genotype calling were performed using ANGSD v0.941 (Korneliussen et al. 2014), employing the GATK genotype likelihood model (-GL 2) and posterior probability-based genotype calling. Missing data per individual was calculated from called genotypes using --missing-indv in VCF tools v0.1.16 (Danecek et al. 2011), and individuals with missing data >30% were excluded from both genotype likelihood and called genotype datasets. Our filtering thresholds reflect common practice for low-coverage whole genome sequencing and were chosen to balance marker retention and data quality (e.g. Lou et al. 2021; Liu et al. 2024). To minimise bias in genetic structure associated with sex-linked scaffolds, we restricted analyses for both datasets to scaffolds with a posterior probability >0.99 of being autosomal in the Galápagos-only dataset (see methods below). See Tables S1 and S2 for further details of filtering.

### Joint inference of sex-linked scaffolds and individual genetic sex

We aimed to infer autosomal scaffolds to avoid inclusion of sex-linked scaffolds that could affect neutral patterns of genetic structure. The sex determination system of any species in the *Philornis* genus remains unknown, however, most dipterans possess an XY sex chromosome system and a well-conserved karyotype (Vicoso and Bachtrog, 2015). Accordingly, we jointly inferred sex-linked scaffolds and the sex karyotype of individuals in the Galápagos-only dataset (prior to filtering of highly related individuals) using BeXY v1.0 (Caduff et al. 2024), which incorporates genotype likelihoods and the *P. downsi* reference genome (Text S3). In parallel, BeXY was used to probabilistically classify scaffolds as autosomal, X-linked, or Y-linked, and to assign genetic sex to individuals (XX, XY, or alternative aneuploidies) based on posterior probabilities. To independently assess confidence in BeXY assigned scaffold classifications, we compared sequencing depth of the inferred autosomes, X- and Y-linked scaffolds between males and females (Text S4).

The *P. downsi* reference genome, derived from a female and not assembled to the chromosomal level (Romine et al. 2022), lacks confirmation of the sex chromosome system. If *P. downsi* follows a standard XY system, Y-linked sequences would be absent from this assembly, precluding reliable identification of Y-linked scaffolds. However, sex determination in Muscidae can deviate from this model; for example, in the house fly (*Musca domestica*), both XY females and XX males can occur, with sex determined by factors on either sex chromosomes or autosomes (Franco et al. 1982; Denholm et al. 1986; Feldmeyer et al. 2007). To avoid potential misclassification due to these uncertainties, putative sex-linked scaffolds were excluded, and all subsequent analyses were restricted to inferred autosomal markers.

### Relatedness, Hardy-Weinberg Equilibrium and Linkage Disequilibrium

To exclude closely related *P. downsi* individuals, pairwise kinship was estimated using relatedness2 in VCFtools v0.1.16 (Danecek et al. 2011) and for pairs with kinship >0.088 (half siblings or closer; Manichaikul et al. 2010), one individual from each pair was removed. Variants were then filtered for Hardy Weinberg Equilibrium (HWE; p >0.001) using ANGSD and linkage disequilibrium (LD; r² <0.5 within 50 kb) using ngsLD v1.2.1 (Fox et al. 2019); the resulting unlinked SNPs were specified for final genotype calling and likelihood estimation in ANGSD.

### Genetic diversity, differentiation and isolation by distance

Genetic diversity and differentiation parameters were estimated separately for the combined dataset (Galápagos and mainland), and for the Galápagos-only dataset. Loma Alta was excluded from genetic diversity analyses due to low sample size. For each island and mainland location, neutral diversity statistics were derived from the Site Allele Frequency (SAF) in ANGSD and the folded site frequency spectrum (SFS) generated using realSFS (Korneliussen et al. 2014). Nucleotide diversity (Θ_π_), Watterson’s theta (Θ_W_) and Tajima’s D were first calculated per scaffold using thetaStat (Korneliussen et al. 2013), and global per-parameter estimates were calculated as the mean across all scaffolds analysed. Global heterozygosity was calculated as the proportion of heterozygous sites from the SFS (Korneliussen et al. 2014).

To calculate pairwise F_st_, individuals from mainland Ecuador (Cerro Blanco, Agua Blanca, Loma Alta) were combined into a single mainland reference group. Global pairwise F_st_ comparisons among the mainland group and each sampled island were calculated from the SAF in ANGSD, with major and minor alleles inferred from genotype likelihoods, and folded joint 2D SFS between groups using realSFS (Korneliussen et al. 2014). We tested for Isolation by Distance (IBD) within the Galápagos dataset using pairwise genetic distances calculated in ngsDist v1.0.10 (Vieira et al. 2016; 1000 bootstraps) and pairwise geographic distances (km) between sampling sites (displayed in Figure 1) using geosphere v1.5.20 (Hijmans 2024) in R. A Mantel test with 999 permutations (vegan v2.7.1; Dixon, 2003) was used to assess the correlation between genetic and geographic distances.

### Genetic structure

Genetic structure was analysed for the Galápagos-only and combined (Galápagos and mainland) datasets separately. Principal component analyses (PCA) were performed using PCAngsd v1.36.1 (Meisner et al. 2021) to generate a covariance matrix. The most likely number of ancestral populations (K) was inferred using both genotype likelihoods with NGSadmix (Skotte et al. 2013) and called genotypes with ADMIXTURE v1.3.0 (Alexander et al. 2009). Both methods were run for K = 1–10. We ran NGSadmix with 10 replicates per K using random seeding, and log-likelihood values were utilised to infer the most likely K. Research demonstrates that assessing model likelihood scores can give a clearer indication of the true K than the ΔK method, which is frequently biased towards K = 2 (Janes et al. 2017) especially when migration is moderate to high (Cullingham et al. 2020). To further validate the most optimal K, ADMIXTURE was run with 10-fold cross-validation (CV) (Alexander et al. 2009). Results of all methods were visualised using ggplot2 (Wickham, 2016) in R v. 4.4.2 (R Core Team, 2024).

### Inferring sex-specific genetic patterns

To assess potential sex-specific genetic dispersal in the Galápagos-only dataset, we conducted pairwise F_st_, IBD and kinship analyses on morphologically determined males and females separately following the methodologies described above. The dataset was split by sex for pairwise F_st_ analyses in ANGSD. For IBD, we partitioned genetic distance matrices by sex and performed Mantel tests separately for males and females. To assess sex-biased relatedness, we compared within-island KING robust kinship estimates (Manichaikul et al. 2010) between males and females using two approaches: (1) calculating mean pairwise kinship within each sex per island, and (2) computing, for each individual, the mean kinship with all other individuals from their island, then averaging these values by sex. This second method accounts for the possibility that sex-biased dispersal may influence the mean kinship of one sex to all other local individuals, not just their own sex (Johnstone and Cant, 2008).

### Effective population size

Contemporary effective population sizes (N_e_) can be estimated using the LD method, however in species with overlapping generations, this estimate reflects the effective number of breeders that produced the sample (N_b_) rather than the effective size for that generation (N_e_) (Waples, 2006). As *P. downsi* exhibits overlapping generations (Causton et al. 2019), we employed the single-sample LD method in NeEstimator2 v2.1 (Do et al. 2014) to estimate N_b_, assuming random mating using the critical values (Pcrit) of 0, 0.01, 0.02 and 0.05 to exclude rare alleles (Waples, 2006). From this estimate, we calculated an adjusted effective population size (N_e (Adj3)_) following the approach of Bilgmann et al. (2021), which corrects for overlapping generations and physical linkage of loci within chromosomes using calculations from Waples et al. (2014, 2016), see Text S5 and Table S3 for details. Prior to N_b_ estimation and to improve SNP independence (Waples et al. 2022; Delord et al. 2025), we used VCFtools --thin to retain one SNP per 10,000 bp in the combined dataset, resulting in a SNP count similar to the Galápagos-only dataset (see Results).

The LD method can provide reliable N_e_ estimates for populations that show genetic isolation (Hill, 1981). Thus, to account for spatial population structure, estimates of N_b_ were calculated among groups of samples according to the genetic clusters we identified across Galápagos and the mainland (see Results). Estimates of N_b_ for each Galápagos island were calculated separately using the Galápagos-only dataset to infer island-level relative differences. As high sampling error can yield negative point estimates and infinite values for confidence intervals (Waples and Do, 2010; Do et al. 2014), NeEstimator2 was run several times for each dataset until an N_b_ estimate could be obtained for every group.

NeEstimator2 includes a bias correction for sample sizes <30, which reduces an inherent upward bias in smaller sample sizes, leading to a slightly downward bias for Pcrit ≥0.05 (Waples and Do, 2010). It is noted that for large census population sizes (N >1000), estimates based on <100 individuals become increasingly unreliable (Waples and Do, 2010). As sample sizes for island and mainland locations are reasonably small considering these thresholds (range: N = 21–161), and considering the bias that can result from low coverage sequencing data (Kardos and Waples, 2024), the purpose of this analysis is to compare relative differences in N_e_ between locations, with estimates of N_e_ interpreted cautiously.

### Directional inter-island migration

To estimate directional migration of *P. downsi* among the Galápagos-only dataset, we employed two approaches: 1) an individual-based, genotype likelihood assignment method (WGSassign v.0.0, DeSaix et al. 2024) and 2) a population-level, called genotype-based migration analysis (divMigrate, Sundqvist et al. 2016). In WGSassign, reference populations were defined by island, and allele frequencies were estimated per reference population using a leave-one-out (LOO) approach. For each individual, reference and assigned Z-scores were calculated from log-likelihoods under each reference population. Reference and assigned Z-scores reflect the confidence of matching to the sampled and highest-likelihood assigned islands, respectively. Individuals assigned with highest likelihood to an island other than their sampled (reference) location were considered candidate migrants. To assess sex-biased migration, we calculated the proportions of male and female candidate migrants Galápagos-wide and within each island, using morphologically determined sex.

The effective sample size (ESS) in WGSassign is a metric estimating the number of fully genotyped individuals needed to match the power of low-coverage sequence data (DeSaix et al. 2024). Since ESS can bias assignment accuracy (DeSaix et al. 2023, 2024), we present standardized ESS of reference populations in WGSassign. Unstandardized ESS was first calculated per island, then the pipeline was re-run using two standardization approaches: (1) iterative random subsampling, and (2) iterative removal of individuals with lowest reference Z-score from the unstandardized analysis. Standardization brought each ESS to within 1.0 of the lowest island ESS. Log-likelihoods and Z-scores were recalculated for each standardized reference population, and candidate migrants were inferred per method. Welch’s unpaired t-tests (Welch, 1947) were used to compare the mean assigned Z-scores among migrants identified by each of the three methods, using the stats package in base R, as the sets of migrants differed between methods.

Directional migration across the Galápagos was inferred at the island level using the divMigrate function (Sundqvist et al. 2016) of the diveRsity v1.9.90 package (Keenan et al. 2013) in R. This approach quantifies relative, directional gene flow between populations based on allele frequencies. The Alcala’s statistic, Nm_Alcala_ (Alcala et al. 2014) was employed to quantify weight and direction of gene flow, which integrates Gst (Nei, 1973) and Jost’s D (Jost, 2008) for robust migration estimates (Sundqvist et al. 2016). Statistical significance of asymmetric migration was assessed with 1000 bootstrap replications (p <0.05) by comparing each population’s genetic distance to a hypothetical migrant pool, distinguishing source (lower distance) from sink (higher distance) populations. The resulting migration network was visualised in R using ggplot2, overlayed to a Galápagos basemap from OpenStreetMap contributors (2026) via NextGIS Data (NextGIS, 2026).

## Results

### Data processing, quality control and scaffold karyotyping

Illumina Novaseq sequencing produced an average of 17,800,301 (range: 11,808,970–74,841,439) paired-end reads per sample. One sample was an outlier, containing almost double the read count of the next highest (38,902,975). To reduce bias from sequence overrepresentation, we down-sampled the outlier by 50% in Picard using random seeding.

Of the 235 samples genotyped in the Galápagos-only dataset, 16 were excluded due to mapping efficacy <50% and 30 were excluded due to >30% missing data that was likely the result of poor DNA quality. We inferred 35 pairs of second-degree relatives (kinship 0.088–0.177; Manichaikul et al. 2010), and to disband these pairs, 20 individuals were excluded. The resulting dataset included 169 individuals with an average depth of coverage 3.27x (range: 2.33–4.86x), and the partitioning of retained individuals across islands, sexes and sampling methods is outlined in Table 1. Of the HWE and LD filtered autosomal dataset, 33,638 SNPs were retained.

**Table 1:** Sample sizes and genetic diversity of *P. downsi* sites across the Galápagos Islands and mainland Ecuador. Global genome-wide estimates of heterozygosity, nucleotide diversity (Θ_π_), Watterson’s theta (Θ_W_) and Tajima’s D among locations are shown (using ANGSD; Korneliussen et al. 2014) for Ecuadorian mainland sites (Cerro Blanco (CB), Agua Blanca (AB) and Loma Alta (LA)) and for Galápagos Islands (Floreana (FL), Isabela (IS), San Cristóbal (SL), Santiago (SO) and Santa Cruz (SZ)). Values are provided from analyses using the combined dataset (Galápagos with mainland Ecuador: 77,973 SNPs), while values in parentheses correspond to the Galápagos-only dataset (33,638 SNPs). N (Total) = number of individuals analysed at each location, partitioned by sex (adult males, adult females, and pupae), and sampling method (trap caught or nest collected). Genetic diversity analyses were performed on the N (Total) for each location.

| Group | Location | N<br>(Total) | N<br>(Male) | N<br>(Female) | N<br>(Pupa) | N<br>(Trapped) | N<br>(Nest) | Heterozygosity | $\Theta_{\pi}$ | $\Theta_w$ | Tajima's D |
| --- | --- | --- | --- | --- | --- | --- | --- | --- | --- | --- | --- |
| Mainland | CB | 16 | 5 | 11 | 0 | 0 | 16 | 0.078 | 0.264 | 0.199 | 0.685 |
|  | AB | 6 | 4 | 2 | 0 | 0 | 6 | 0.190 | 0.306 | 0.255 | 0.554 |
|  | LA* | 2 | 1 | 1 | 0 | 0 | 2 | - | - | - | - |
| Galápagos | FL | 25 (27) | 11(12) | 14(15) | 0(0) | 25(27) | 0(0) | 0.021 (0.019) | 0.250 (0.331) | 0.156 (0.201) | 1.150 (1.298) |
|  | IS | 52 (54) | 29(29) | 23(25) | 0(0) | 11(11) | 41(43) | 0.006 (0.004) | 0.240 (0.320) | 0.131 (0.170) | 1.372 (1.539) |
|  | SL | 20 (21) | 9(10) | 10(10) | 1(1) | 3(4) | 17(17) | 0.030 (0.033) | 0.232 (0.310) | 0.145 (0.190) | 1.157 (1.272) |
|  | SO | 33 (33) | 12(12) | 21(21) | 0(0) | 33(33) | 0(0) | 0.012 (0.009) | 0.248 (0.331) | 0.148 (0.193) | 1.226 (1.373) |
|  | SZ | 31 (34) | 10(11) | 21(23) | 0(0) | 31(34) | 0(0) | 0.015 (0.010) | 0.250 (0.331) | 0.151 (0.193) | 1.175 (1.350) |

\* Omitted from genetic diversity analysis due to low sample size.

All 39 mainland samples in the combined dataset had mapping efficacy >50%. Missing data percentage per individual was calculated across the mainland and Galápagos for the combined dataset, and 10 individuals (8 Galápagos; 2 mainland) were excluded due to >30% missing data. Kinship assessment inferred 14 pairs of second-degree relatives among the mainland samples, all within nests, and 13 individuals were excluded to disband pairs. The resulting combined dataset included 185 individuals across Galápagos (161) and the mainland (24) with an average depth of coverage 3.57x (2.33–7.56x). After filtering for autosomes, HWE and LD, a total of 77,973 SNPs were identified in the combined dataset.

### Classification of scaffolds and assignment of individual genetic sex

An XX/XY karyotype system was inferred from the dataset, consistent with the most common sex determination system for dipterans (Vicoso and Bachtrog, 2015). Of the 32,922 scaffolds analysed, posterior probabilities inferred 28,073 autosomal scaffolds (85.3%), 2,656 X-linked scaffolds (8.1%), and 2,193 Y-linked scaffolds (6.7%) (Figure S2a). Most scaffold assignments were made with high confidence (Figures S2b–d), however we interpret the putative Y-linked scaffolds with caution, as their identification may reflect misassembled or complex genomic regions resulting from the use of a female-derived reference genome. A total of 27,926 scaffolds were inferred as autosomal with >99% confidence and were retained in the dataset (Figure S2b).

Across the Galápagos dataset (prior to filtering for genetic relatedness), 134 individuals were identified as XX and 54 individuals were identified as XY (Figure S3a). All individuals were assigned karyotypes with a posterior probability of 1.0 except for a single morphological female from SL assigned as XX with posterior probability 0.64 (Figure S3b,c). Genetic sex assignments did not always align with morphological identification of sex, likely due to uncertainty inherent within the estimations due to data resolution, a lack of sex karyotype knowledge for *P. downsi,* and a female-derived reference genome. Of the 74 morphologically determined Galápagos males, 32 (43%) were assigned as XX, while of the 94 morphological females, three (0.03%) were assigned as XY. One morphological female was assigned a haploid (X) sex karyotype, likely due to high missing data (28%) and low read depth (2.3x) resulting in insufficient coverage for confident XX or XY assignment. The dataset included one pupa assigned as XX, which could not be sexed, and was therefore labelled by genetic sex in our PCA but excluded from sex-specific analyses.

Autosomal scaffolds showed a depth ratio of 1.05 between morphologically determined sexes and a ratio of 1.07 between XY and XX assignments (Figure S4), closely matching the expected 1:1 ratio for an XY system (Palmer et al. 2019). Therefore, we retained putative autosomal scaffolds for further analyses. Depth ratios for the inferred sex-linked scaffolds generally followed the expected XY patterns (Palmer et al. 2019). However, some Y-linked sequence reads were detected in females, and X-linked scaffolds had higher than expected depth in males (Figure S4). Putative sex-linked scaffolds were not included in subsequent analyses.

### Genetic diversity, differentiation and isolation by distance

Mainland Ecuador locations (n = 2) exhibited higher heterozygosity (CB: 0.078; AB: 0.190), nucleotide diversity (Θ_π_) (CB: 0.264; AB: 0.306), genetic diversity as measured by Watterson’s theta (Θ_W_) (CB: 0.199; AB: 0.255), but lower Tajima’s D (CB: 0.685; AB: 0.554) compared to the five sampled Galápagos islands pooled (Table 1). For the combined SNP datasets (five islands and mainland) and Galápagos-only dataset (n = 5 islands), SL *P. downsi* showed the greatest heterozygosity of the Galápagos sites (Combined: 0.030; Galápagos-only 0.033) and IS showed the lowest heterozygosity (Combined: 0.006; Galápagos-only: 0.004) (Table 1). All Tajima’s D values were positive, indicating a deficit of rare alleles, suggesting balancing selection or population contraction (Schmidt and Pool, 2002). For the Galápagos-only dataset, the lowest Tajima’s D was observed on SL (1.272), and on FL for the combined dataset (1.150), with SL close behind (1.157). The highest Tajima’s D was observed on IS for both datasets (Combined: 1.372; Galápagos-only: 1.539). Nucleotide diversity (Galápagos-only range: 0.232–0.250; combined range: 0.310–0.331) and Watterson’s theta (Galápagos-only range: 0.131–0.156; combined range: 0.170–0.201) were similar across islands. For both datasets, nucleotide diversity was lowest on SL and highest (and equal) on FL and SZ, and Watterson’s theta was lowest on IS and highest on FL (Table 1).

The greatest genetic differentiation occurred between the mainland reference group (CB, AB, LA) and each of the Galápagos islands (mean F_st_ = 0.180, range: 0.172–0.197), with SL showing the highest differentiation from the mainland (0.197) while the other islands were similarly differentiated from the mainland (mean F_st_ = 0.176; range: 0.172–0.179; Figure S5). Among the Galápagos islands, SL was consistently the most differentiated (combined dataset mean F_st_ = 0.057, range: 0.053–0.062, Figure S5; Galápagos-only mean F_st_ = 0.051, range: 0.048–0.057, Figure S6). In contrast, lower genetic differentiation was observed between FL, IS, SZ and SO (combined dataset mean F_st_ = 0.014, range: 0.007–0.018, Figure S5; Galápagos-only mean F_st_ = 0.012, range: 0.005–0.015, Figure S6), with SZ and SO being the most genetically similar in both datasets (combined dataset mean F_st_ = 0.007, Figure S5; Galápagos-only mean F_st_ = 0.005, Figure S6). A significant but weak IBD relationship was detected across the Galápagos Islands (Mantel r = 0.207, CI = 0.174–0.230, p < 0.001; Figure S7).

### Genetic structure

In the combined dataset, PCA revealed clear genetic divergence between the Galápagos and the mainland, with SL individuals clustering separately from the other four islands; PC1 and PC2 explained 10.8% and 3.9% of the genetic variation, respectively (Figure 2a). The NGSadmix log-likelihoods showed best support for K = 2, with further subdivision at K = 5 (Figure S8a). At K = 2, admixture coefficients revealed complete divergence between Galápagos and mainland (Figure 2b). The lowest CV error in ADMIXTURE best supported K = 5 (Figure S8b), with mainland samples forming one cluster, while Galápagos Islands did not group strictly by island and exhibited mixed ancestry (Figure S9a).

**Figure 2:**
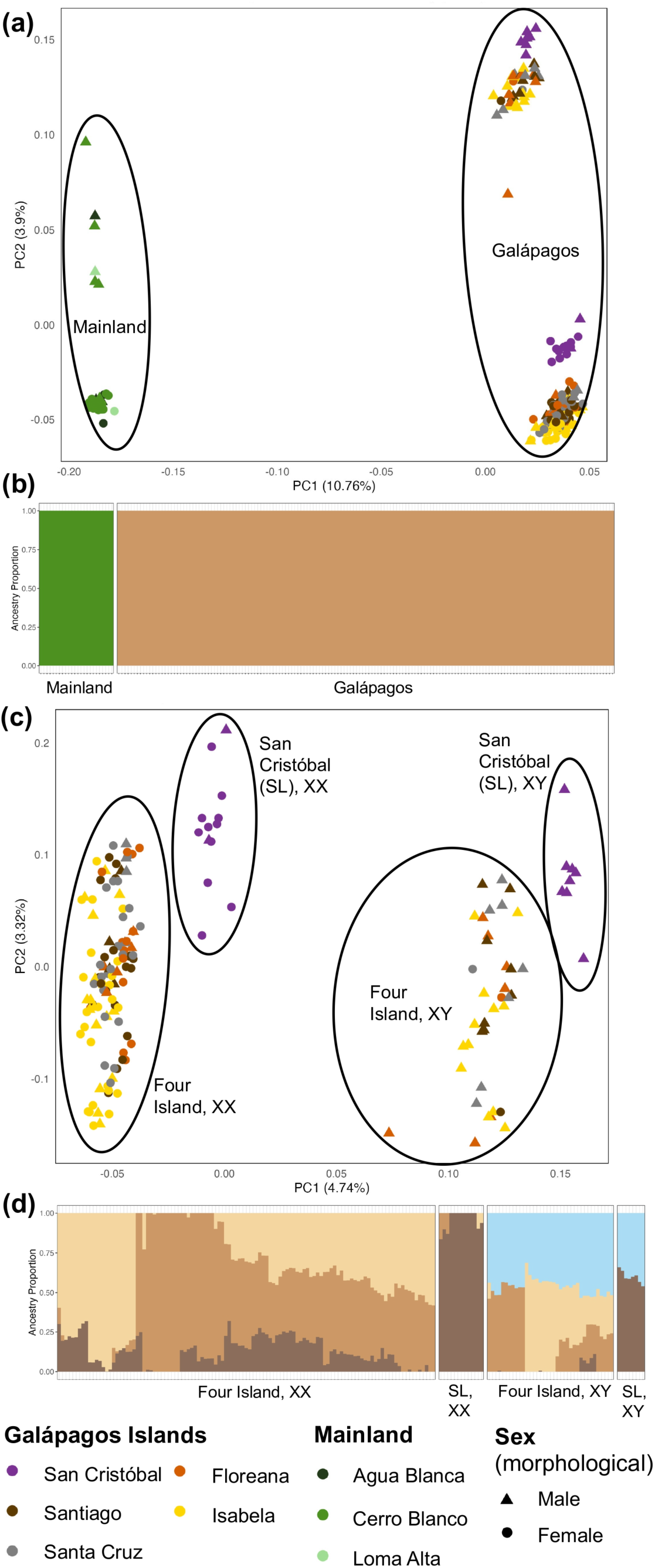
Neutral, putative autosomal population structure of *P. downsi* across the Galápagos archipelago and mainland Ecuador. a) Principal components analysis (PCA), calculated with PCAngsd (Meisner et al. 2021), depicts genetic separation between Galápagos and mainland, b) NGSadmix (Skotte et al. 2013) analysis of Galápagos with mainland best supports K = 2, separating Galápagos and mainland into distinct clusters. c) PCA for Galápagos-only dataset shows greatest separation between XX and XY groups, identified with BeXY (Caduff et al. 2024), despite using a putative autosomal dataset. Divergence is also evident between San Cristóbal (SL) and the other islands (Four Island). d) NGSadmix analysis of Galápagos-only dataset best supports K = 4. Islands are admixed and clusters do not correspond to locations, but genetic sex assignment shows corresponding clusters across each group. Point symbols represent the morphologically determined sex, which varied from their inferred genetic sex.

In the Galápagos-only dataset, two main genetic clusters were present with two other, less diverged, sub-clusters evident within the PCA, with these subclusters separated predominantly as XX and XY genetic sexes (Figure 2c). Notably, SL samples formed discrete XX and XY groups that were divergent from the four other islands (Figure 2c). The IS individual that was putatively assigned as X was grouped with the XX cluster across the four other islands. Notably, the percentage of genetic variation explained by the first two PC axes was small (PC1 = 4.7% and PC2 = 3.3%; Figure 2c). The PCA clusters were not attributable to sequencing batch, missing data, or estimated depth biases (Figure S10).

For the Galápagos-only dataset, the most likely number of ancestral populations was identified as K = 4 using both log-likelihoods and CV error (Figure S11), however as for the PCA above, this did not entirely correspond with island location, and was largely explained by genetic sex in addition to genetic divergence of SL. Ancestral clusters inferred from genotype likelihoods (Figure 2d) and called genotypes (Figure S9b) were broadly consistent for K = 4. These clusters did not entirely correspond with geographic sampling locations, except that one cluster was more common on SL, while two clusters were more common across the four other islands (Figure 2d). As genetic sex-biased structure was observed in our PCA, we examined the alignment between ancestry patterns and the genetic sex of individuals. We found that one cluster was exclusive to individuals putatively assigned as XY for both SL and the other four islands (Figure 2d).

### Sex-biased genetic structure

No evidence for sex-biased genetic structure was detected via IBD, as both females (r = 0.173, CI: 0.140–0.197, p = 0.001) and males (r = 0.177, CI: 0.140–0.208, p = 0.001) showed similarly weak but significant IBD (Figure S12). Global pairwise F_st_ values were also comparable between males and females across islands (Figure S13). Within islands, mean pairwise kinship differed by only 0.001 between sexes (range: 0.0–0.002; Table S4), and relative to the total sample, the difference averaged 0.0004 (range: 0.0–0.001; Table S5). Thus, relatedness levels were similar for males and females on each island, and no signature of sex-biased genetic dispersal was evident.

### Effective population size

The raw N_b_ estimates increased slightly as the Pcrit threshold was lowered, but relative differences among locations persisted (Table S6). As Pcrit ≥0.05 reduces sampling bias in smaller population sizes (Waples and Do, 2010), results for this threshold are presented. Raw N_b_ estimates were low across locations (range: 117–842; Table S6) and decreased slightly after adjustment for overlapping generations (N_b(Adj2)_ range: 105–763; Table S7). The derived N_e_ from this adjustment slightly increased estimates (N_e(Adj2)_ range: 218–1,575; Table S8), while correction for chromosome number yielded substantially higher estimates (N_e(Adj3)_ range: 2,229–16,073; Table S9).

The pooled mainland N_e(Adj3)_ (11,921, CI = 6,038–268,866) was considerably higher than the five island Galápagos group (2,229, CI = 2,032–2,468), though with greater uncertainty (Table 2). Among pooled Galápagos groups, the four island N_e(Adj3)_ (3,110, CI = 2,891–3,360) exceeded the five island estimate – possibly due to dataset differences or inclusion of genetically divergent SL in the five island group. Within the two genetic clusters of the Galápagos-only dataset, the four-island cluster had a lower N_e(Adj3)_ than SL (3,623, CI= 1,981–17,441; Table 2). Individual island N_e(Adj3)_ estimates were broadly similar with overlapping confidence intervals. Estimated N_e(Adj3)_ for SL was intermediate to other islands, but with a substantially higher upper confidence limit (Table 2). The IS N_e(Adj3)_ was lowest (2,260, CI = 2181–3373), while FL (3,801, CI: 2,816–5,773), SO (4,079, CI: 3,166–5,680) and SZ (4,283, CI: 3,352–5,886) were similar (Table 2).

**Table 2:** Effective number of breeders in sample (N_b_) and bias-corrected effective population size (N_e(Adj3)_) of *P. downsi* across the Galápagos and mainland Ecuador. The N_b_ was estimated with the Linkage Disequilibrium (LD) method in NeEstimator2 (Do et al. 2014) using the critical value of 0.05 and 95% JackKnife confidence intervals (CI). The N_e(Adj3)_ was derived from N_b_, correcting for overlapping generations and physical linkage of loci within chromosomes. The five-island Galápagos group includes all islands (Isabela, Floreana, Santiago, Santa Cruz, San Cristóbal), while the four-island group excludes San Cristóbal, a genetically divergent island.

| Dataset | Location | $N_b$ | CI ($N_b$) | | $N_{e(Adj3)}$ | CI ($N_{e(Adj3)}$) | |
| --- | --- | --- | --- | --- | --- | --- | --- |
|  |  |  | L | U |  | L | U |
| Galápagos-only | Isabela<br>(N = 54; 33,440 SNPs) | 139 | 114 | 177 | 2660 | 2181 | 3373 |
|  | Floreana<br>(N = 27; 33,072 SNPs) | 199 | 147 | 302 | 3801 | 2816 | 5773 |
|  | Santiago<br>(N = 33; 33,373 SNPs) | 214 | 166 | 298 | 4079 | 3166 | 5680 |
|  | Santa Cruz<br>(N = 34; 33,367 SNPs) | 224 | 176 | 308 | 4283 | 3352 | 5886 |
|  | San Cristóbal<br>(N = 21; 29,025 SNPs) | 190 | 104 | 914 | 3623 | 1981 | 17441 |
|  | Four Island<br>(N = 148; 33,587 SNPs) | 163 | 151 | 176 | 3110 | 2891 | 3360 |
| Galápagos and mainland | Galápagos (five islands)<br>(N = 161; 22,940 SNPs) | 117 | 106 | 129 | 2229 | 2032 | 2458 |
|  | Mainland<br>(N = 24; 27,883 SNPs) | 625 | 316 | 14095 | 11921 | 6038 | 268866 |

### Directional inter-island migration

The WGSassign analysis found different sets of candidate migrants depending on the method: 34 in the unstandardized analysis (mean assigned Z-score: 20.50; range: 8.15–34.84), 40 using reference ESS standardized by random subsampling (mean: 13.81; range: -10.00–34.84), and 46 using standardization by subsampling highest reference Z-scores (mean: 13.40; range: -8.25–34.84) (Figure 3). Eighteen candidate migrants overlapped among all methods (Figure S14). Mean assigned Z-scores of candidate migrants were statistically higher in the unstandardized method than with random subsampling (t = 3.25; df = 62.00; p = 0.002) or highest Z-score subsampling (t = 3.77; df = 74.04; p < 0.001), while the two standardization methods did not differ (t = 0.17; df = 80.90; p = 0.86). The unstandardized method also produced fewer candidate migrants and likely offered the most accurate assessment of inter-island movement. Across all three methods, most individuals were assigned to their sampled island (Figure 3).

**Figure 3.**
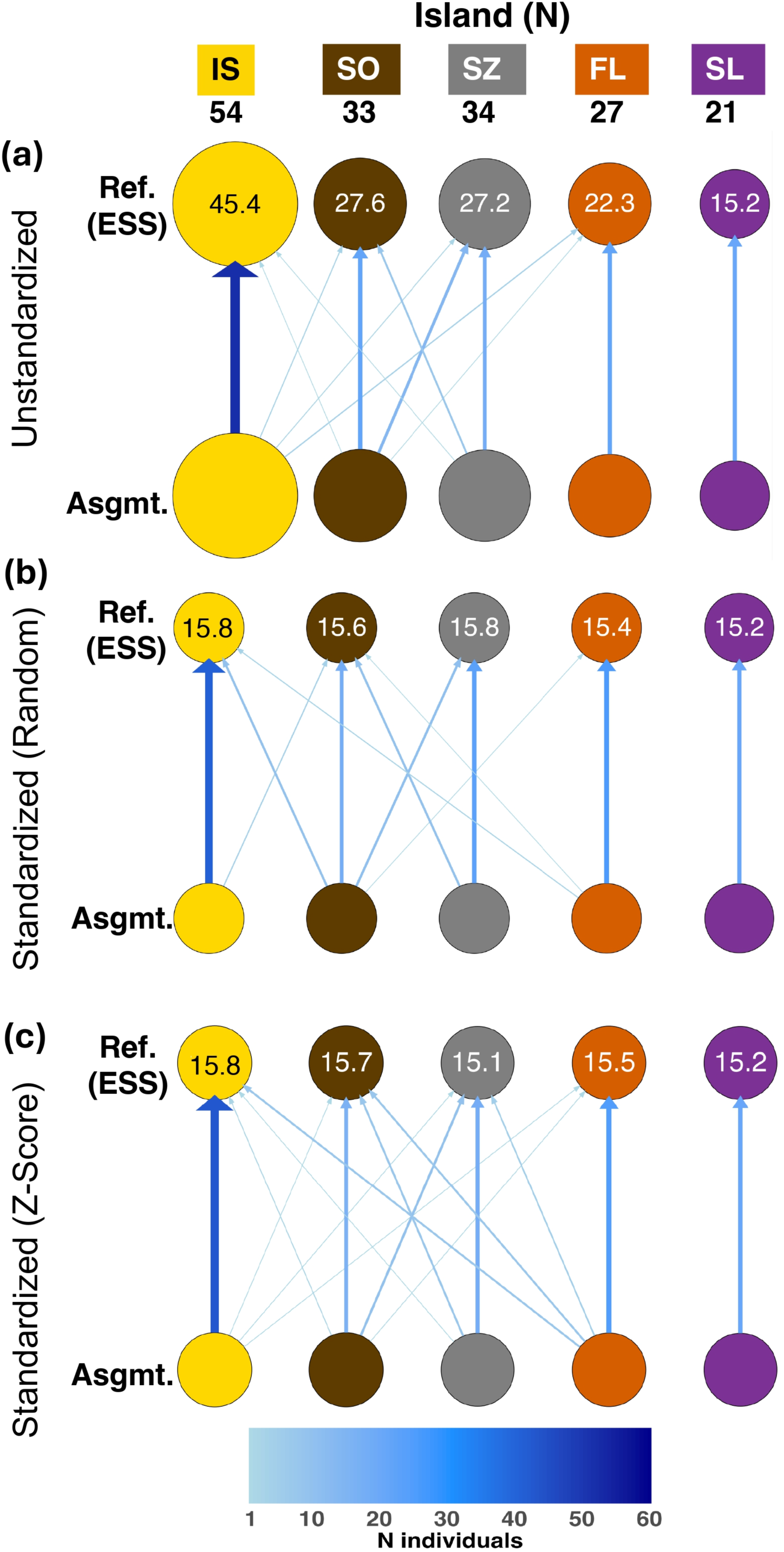
Population assignment of *P. downsi* individuals among Galápagos Islands using WGSassign (DeSaix et al. 2024). Coloured circles represent islands: yellow = Isabela (IS), brown = Santiago (SO), grey = Santa Cruz (SZ), orange = Floreana (FL), purple = San Cristóbal (SL). Total sample size is indicated below island labels. Within each panel, the upper circle indicates the island where individuals were sampled from (reference population, “Ref.”), and the lower circle indicates the putative island of origin (assigned population, “Asgmt.”). Circle size is proportional to the reference population effective sample size (ESS). Arrow colour and thickness scales with the number of assigned individuals. Panels compare methods for standardizing ESS in reference populations: (a) no ESS standardization; (b) ESS standardized via random subsampling per island; (c) ESS standardized by subsampling individuals with highest reference Z-scores.

No candidate migrants were associated with SL, inferring greatest isolation (Figure 3). Islands SZ and SO consistently emerged as main inter-island dispersal hubs, exchanging the most migrants with each other and other islands (Figure 3). Island IS showed higher connectivity than FL, with numbers of migrants varying by method (Figure 3). No evidence for sex-biased migration was found across assignment methods. In the unstandardized analysis, similar proportions of females (19/94; 20.2%) and males (15/74; 20.3%) were assigned to non-reference islands. Although sex-biased proportions varied somewhat by island, migrant numbers for each sex were too low to detect any clear sex-biased signal (Table S10).

Relative migration analyses (divMigrate) identified SL as the most isolated island, with significant asymmetric gene flow from SL to other islands (range: 0.27–0.34) exceeding return gene flow (range: 0.14–0.16); no other island showed significant asymmetry (Figure 4). Relative migration patterns mirrored inter-island F_st_ results, with the highest gene flow between SZ and SO (1.0 SZ→SO, 0.99 SO→SZ), indicating strong connectivity (Table S11). High connectivity was also observed between IS with SO (0.85 IS→SO; 0.88 SO→IS) and IS with SZ (0.74 IS→SZ; 0.77 SZ→IS). FL displayed moderate connectivity, which was greatest with IS (0.70 FL→IS; 0.63 IS→FL), followed by SO (0.65 FL→SO; 0.61 SO→FL), then SZ (0.59 FL→SZ; 0.56 SZ→FL) (Table S11).

**Figure 4:**
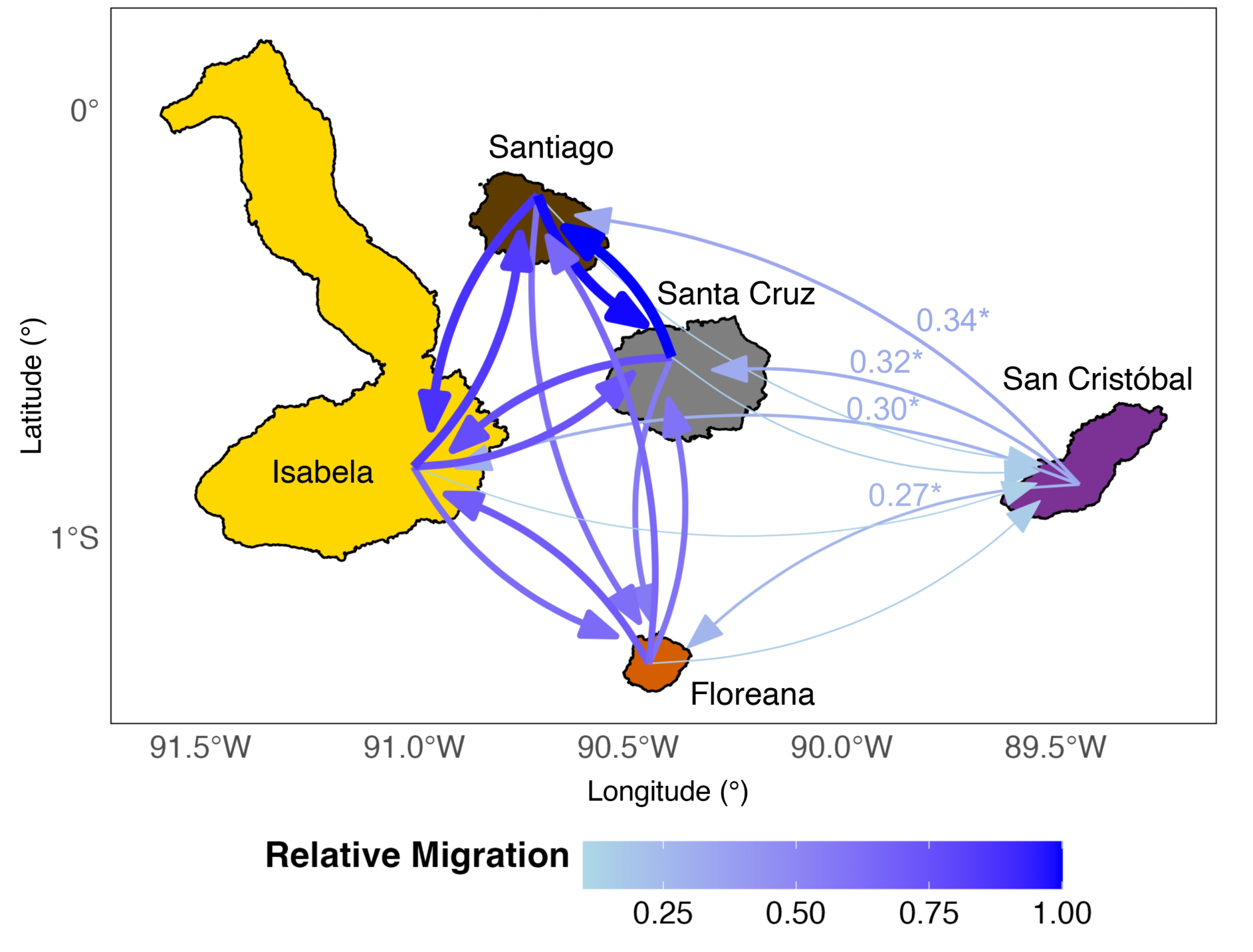
Relative directional gene flow of *Philornis downsi* across the Galápagos archipelago, estimated using divMigrate (Sundqvist et al. 2016). Migration rates between islands were calculated under the *Nm* model (Alcala et al. 2014). Arrow colour and thickness represent relative migration rates. The position of origin and termination of arrows represents general inter-island migration between islands, not to exact sampling localities. Significant values of asymmetric migration are indicated next to corresponding arrows with asterisks, and in this analysis, only asymmetric migration from San Cristóbal to all other islands is significant.

## Discussion

Here we present a comprehensive analysis of genome-wide genetic diversity and structure in the invasive avian ectoparasitic fly, *Philornis downsi,* with sampling across the Galápagos Islands and western mainland Ecuador. Inter-island gene flow was high among islands except for San Cristóbal (SL), which showed distinct genetic divergence, the highest heterozygosity, significant asymmetric genetic migration, and a much higher upper confidence limit for effective population size than other islands. These patterns are consistent with a source-sink dynamic where multiple founders first invade San Cristóbal before westward dispersal. Despite weak sex-associated genetic structure, no evidence was found for sex-biased dispersal. Strong genetic divergence and genetic clustering between island and mainland *P. downsi* indicates a founder effect within the Galápagos (consistent with Basnet et al. 2025). This is the first study to reveal clear genetic structure and asymmetric genetic migration in *P. downsi* across the Galápagos archipelago, with implications for island-specific fly control strategies.

### Genetic structure and genetic diversity across islands

Consistent with prior studies (Dudaniec et al. 2008; Koop et al. 2021; Common et al. 2023; Basnet et al. 2025), most islands showed high gene flow and low genetic differentiation, mainly corresponding with inter-island geographic distances. Closely situated islands – Isabela, Santa Cruz, and Santiago (16–27 km apart) – had the highest gene flow, and notably, Isabela had the lowest genetic diversity despite being the largest island. Floreana (50 km south of Santa Cruz) showed intermediate gene flow and genetic diversity. San Cristóbal was the most genetically distinct and genetically diverse island, and the most geographically isolated (66 km southeast of Santa Cruz). Basnet et al. (2025) also found their most isolated island (Marchena) was distinctly structured, but elevated diversity was not reported. While the small San Cristóbal sample in Koop et al. (2021) (N = 5) did not cluster independently or show greater diversity, it displayed higher F_st_ from the other islands, supporting our findings with a larger sample (N = 21). Overall, this combination of high isolation, differentiation, and diversity suggests that San Cristóbal may harbor a larger, more stable *P. downsi* population or have experienced a unique colonization history.

Effective population sizes further contextualize patterns of diversity and structure. San Cristóbal showed moderate N_e_ but a high upper confidence limit, suggesting that its true N_e_ may be much higher than our data could accurately estimate – consistent with its increased genetic diversity. Notably, San Cristóbal had the smallest sample size, which originated mostly from nests within a small geographic area, possibly underestimating genetic diversity. Isabela had the lowest N_e_ and genetic diversity, supporting a genetic bottleneck. However, all islands showed overlapping confidence intervals, and broader sampling is needed for more accurate N_e_ estimates.

Although we corrected N_e_ for overlapping generations and linkage, *P. downsi’s* life history traits – fluctuating population sizes and sex ratios (Causton et al. 2019) – can greatly reduce N_e_, especially in highly fecund insects (Frankham 1995). Marked declines in abundance during the host interbreeding season (Bulgarella et al. 2022) likely result in N_e_ estimates far lower than census sizes. Further, gene flow across islands violates assumptions of isolated, randomly mating populations (Waples, 2006; Kardos and Waples, 2024). Thus, island-specific N_e_ values are interpreted cautiously and only as relative comparisons.

Human-mediated introductions may contribute to the genetic diversity patterns observed, particularly on San Cristóbal. Multiple introductions can increase genetic diversity in invasive species (Novak and Mack, 1993; Dlugosch and Parker, 2008), and San Cristóbal – being closest to the mainland and the first docking point for cargo ships from Ecuador (Bigue et al. 2012; Ministerio de Transporte y Obras Pública 2022) – may serve as the main entry point for *P. downsi* to enter the Galápagos. Imports of fruit and vegetables to San Cristóbal, which are part of the adult fly’s diet (Fessl et al. 2018), could facilitate introductions. Notably, genetic evidence from the introduced tree, *Cedrela odorata*, similarly points to San Cristóbal and/or Santa Cruz as primary introduction sites before spreading to other islands (Albuja-Quintana et al., 2024). In contrast, Isabela is the most distant and the last island on the major cargo shipping route (Bigue et al. 2012; Ministerio de Transporte y Obras Pública 2022), supporting its role as a potential sink. If *P. downsi* enters San Cristóbal from cargo ships arriving from the major city of Guayaquil, one might expect Cerro Blanco to be a source population given its proximity to Guayaquil (15km NW from Port of Guayaquil), but we found no evidence for this. Plausibly, migrant *P. downsi* may board ships in Guayaquil from other mainland sources, thus a genomic analysis of flies from South American regions exporting produce to Guayaquil could help identify introduction origins.

### Asymmetric westward migration from San Cristóbal Island

Asymmetric migration patterns further support San Cristóbal as a source population. Higher outgoing than incoming gene flow may arise from either ongoing contemporary migration or a strong and recent founder effect, as source populations often retain higher genetic diversity (Sundqvist et al. 2016). Demographic analysis is needed to identify mainland sources and clarify whether these patterns reflect ongoing migration or demographic history. Nevertheless, multiple lines of evidence support San Cristóbal as a migrant source: elevated genetic diversity, potentially higher N_e_, increased genetic differentiation, and asymmetric westward migration collectively suggest either ongoing westward dispersal after multiple mainland introductions, or a historical role as the initial colonization point for *P. downsi*. Additionally, Isabela’s genetic bottleneck may result from limited gene flow from San Cristóbal, which is both geographically and genetically the most distant island from Isabela.

Natural dispersal mechanisms, such as wind, likely interact with human-mediated transport to shape genetic migration patterns. Wind-assisted dispersal is well documented in dipterans (Hogsette and Ruff, 1985; Welty Peachey et al. 2025; Darnis et al. 2025), including aerial movement across the Galápagos (Peck 1994), and Muscidae exhibit transoceanic dispersal capabilities (Holzapfel and Harrell, 1968; Löwenberg-Neto and de Carvalho, 2020). Predominant southeasterly trade winds transport air from South America northwest across the archipelago (Cinay et al. 2025; Forryan et al. 2021; Trueman and d’Ozouville, 2010), positioning San Cristóbal as the eastern gateway for migrants dispersed westward. Although we find no genetic structure in *P. downsi* within islands, consistent with Dudaniec et al. (2008), the fly’s inter-island flight capacity remains undocumented (Fessl et al. 2018).

In addition to wind-assisted movement and active flight, insect dispersal across the Galápagos is likely influenced by accidental human transport (Roque-Albelo and Causton, 1999). Tourism and residential growth over the last 25 years has increased inter-island maritime traffic (Denkinger and Vinueza, 2014; Alava et al. 2023), and multiple insects orders have been documented travelling between Galápagos islands on ships (Silberglied 1978). Adult *P. downsi* are likely transported between islands via boats (Fessl et al. 2018), with documented instances of individuals ‘hitchhiking’ on tourist vessels (Lomas 2008). Santa Cruz, the most populous island and main hub for inter-island boat travel (Parra et al. 2013; Keith et al. 2016), likely serves as a hub for *P. downsi* dispersal due to its central geographic position and high traffic, facilitating symmetric short-distance gene flow between neighbouring islands through boat hitchhiking and flight. Conversely, longer-distance asymmetric gene flow from San Cristóbal appears assisted by southeast trade winds, with boat transport as an additional vector of transport.

### Genetic differentiation of native mainland and invasive island populations

Our findings corroborate studies reporting founder effects and genetic bottlenecks in Galápagos *P. downsi* (Dudaniec et al. 2008; Koop et al. 2021; Common et al. 2023; Basnet et al. 2025). Substantially lower N_e_ and genetic diversity on the Galápagos compared to mainland Ecuador supports a founder effect for invasive island populations. Positive Tajima’s D across all groups, with higher values in the Galápagos, is consistent with a recent genetic bottleneck (Schmidt and Pool, 2002; Ramakrishnan et al. 2005). Elevated genetic diversity on San Cristóbal suggests ongoing introductions, while evidence of genetic admixture in the Galápagos range suggests either colonization by different ancestral lineages or post-introduction genetic divergence. As Galápagos and the mainland form discrete ancestral clusters, our mainland sites – also sampled by Koop et al. (2021) and Basnet et al. (2025) – are unlikely to represent source populations for Galápagos invasion, as also stated by Basnet et al. (2025).

### Determining sex via genetic estimation

Putative genetic-based sex assignments did not always match morphological identifications, despite adult *P. downsi* being sexually dimorphic and reliably sexed using morphology (Dodge and Aitken, 1968; Fessl et al. 2001; Lahuatte et al. 2025). We are confident in our field identifications, supported by high-resolution morphological images (Figure S1), so discrepancies are unlikely due to misidentification. While dipterans typically have XY sex chromosomes, their sex determination system can be variable, even within a species (Vicoso and Bachtrog, 2015). For example, in house flies (*Musca domestica*), polygenic and autosomal sex determination can emerge (Hiroyoshi et al. 1982; Hamm et al. 2015; Meisel et al. 2016), resulting in population-level variation and diverse karyotypes, including frequent XX males (Denholm et al. 1986; Cakir and Kence, 1996; Feldmeyer et al. 2007). We inferred a high proportion of homogametic males and fewer heterogametic females, suggesting either mixed/non-standard sex determination in *P. downsi,* or limitations in identifying males. As BeXY (Caduff et al. 2024) infers genetic sex from sex chromosome ploidy, the absence of Y-linked scaffolds in our alignment to a female *P. downsi* reference genome, which was also somewhat fragmented (37,222 scaffolds; Romine et al. 2021), together with the limited sequencing depth of our data, may have hindered reliable XY male classification. Chromosome-level reference genomes for both sexes are needed to clarify sex determination in *P. downsi*.

### Sex-associated genetic structure

Sex-associated genetic structure in *P. downsi* was weak but consistent and could not be explained by technical or sequencing biases. This pattern matches the sex-biased ‘double banded’ clustering found by Basnet et al. (2025) in the Galápagos islands but not in the mainland (S. Lamichhaney, personal communication). Our sex-biased clustering aligned with genetic sex assignments and, as analyses focused on autosomes, suggests autosomal sex determination rather than classic sex-linked divergence (e.g. Loeschcke et al. 1993, Schütt and Nöthiger, 2000, Hamm et al. 2015). Some residual sex-chromosome signal may persist, given observed coverage deviations for an XY system (Figure S3) and possible absence of Y-linked scaffolds in the female reference genome (Romine et al. 2022). No evidence for sex-biased genetic migration was detected, so dispersal is an unlikely cause, though larger sample sizes or demographic approaches may provide further clarity. Sex-specific selection may also contribute, and has previously been documented morphologically in *P. downsi*, whereby downward fecundity selection on female abdomen size has occurred over recent decades (Common et al. 2020). We interpret the weak sex-associated sub-structure cautiously given high genetic admixture in our Galápagos dataset, which can affect PCA clustering interpretation (McVean 2009).

### Management implications

Our findings inform management of invasive avian parasitic flies on the Galápagos, where current control relies on manual insecticide applications and self-fumigating nesting materials (Knutie et al. 2014; Kofler et al. 2025b). The host-specific parasitoid wasp *Conura annulifera* is a promising biological control agent (Boulton and Heimpel, 2017; Bulgarella et al. 2017; Heimpel, 2017; Boulton et al. 2019; Ramirez et al. 2025, 2022). Effective invasive species management on archipelagos depends on low reinvasion risk (Pichlmueller et al. 2020; Kumar et al. 2022), and population genetic information can guide biological control releases. Resistance to biological control agents is more likely to evolve in genetically diverse populations (Hufbauer and Roderick, 2005), whereby subsequent invaders encounter novel environmental conditions that trigger new adaptive genetic processes that could dampen control effectiveness (Banks et al. 2018). Therefore, from a genetics perspective, isolated islands are preferred sites for biological control due to lower numbers of new arrivals and lower gene flow. Although San Cristóbal was the most genetically isolated island, its heightened genetic diversity and possible migrant influx from unsampled mainland areas suggest control here may require a multi-faceted strategy with enhanced biosecurity measures. By contrast, Floreana, with intermediate genetic diversity and inter-island gene flow compared to other islands, may offer a more efficient target. Ultimately, biological control actions must also consider island access, infrastructure and community involvement alongside population genomic insights.

## Conclusion

Our study reveals genetic structure, asymmetric inter-island migration, and differences in genetic diversity between invasive and native ranges of *P. downsi* across the Galápagos islands. San Cristóbal emerges as a likely point of entry and a source population for westward dispersal, highlighting the interplay between geography, human-mediated transport, and natural dispersal processes in shaping invasion dynamics. While weak sex-biased genetic structure was present when analysing putative autosomal markers only, it was unexplained by sex-biased dispersal. Our findings show that island-specific control strategies for *P. downsi* can benefit from understanding gene flow, directional genetic migration and effective population size across the islands. More broadly, we demonstrate how integrating population genomics with ecological and anthropogenic context can illuminate the processes driving biological invasions in island systems, providing an example for predicting and mitigating impacts of invasive species elsewhere.

## Acknowledgments

We thank the Galápagos National Park Directorate (GNPD) and the Ecuadorian Institute of Biodiversity (INABIO) for granting permission to conduct this study (PC-02-20, PC-73-21, PC-33-24, PC-22-23 and Contrato Marco MAE-DNB-CM-2016-0043, MAE-DNB-CM-2016-0045, MAAE-DNB-CM-2020-0133, MAAE-CMARG-2021-0319 and MAATE-DBI-CM-2024-0358). This project was funded by the Austrian Science Fund (FWF) [10.55776/PAT1115224], [10.55776/P36342] and [10.55776/W1262], the Galápagos Conservation Trust Fund, Macquarie University funding [to A Hay], the International Community Foundation (with a grant awarded by the Leona M. and Harry B. Helmsley Charitable Trust), the National Geographic Society, Galápagos Conservancy, the University of Minnesota, and the National Science Foundation [DEB-1949858 to S A Knutie]. For logistical support we thank the staff at the Charles Darwin Research Station (CDRS), especially Marta Romoleroux, Nicolas Padilla, and the Galápagos Science Center. We thank Eric Horstman at the Fundacion pro-Bosque for facilitating our research at the Cerro Blanco Protected Forest, and the Rosales family for logistical support at Loma Alta. We thank Paolo Piedrahita, Frecia Pinguil and Maria Beatriz Perez for logistical support at ESPOL University in Guayaquil, and Jefferson García-Loor for his incredible logistical support for the entire field work operation in Galápagos. We thank Katherine Anabel Albán Morales, Gabriel Brito, Tim Garret, Agustín Gutierrez, Alena Hohl, Leon Hohl, Melanie Kaluppa, Johanna Kniely, Matthew Leaf-Milham, Caelan Linke, Denis Mosquera, Lorena Rojas Allieri, Elizabeth Tituaña Gil, Mauricio Torres, Isabela Vargas and Fernando Villegas for their help in the field. We thank our guides and porters Novarino Castillo Gonzaga, Alfredo Abad Castillo and Carlos Salinas for their support and guidance on Santiago Island. We thank the communities on all inhabited islands for their warmth and kindness. This publication is contribution number [TBD] of the Charles Darwin Foundation of the Galápagos Islands.

## Data Accessibility and Benefit Sharing

<u>Data Accessibility:</u> Raw sequence data will be uploaded to the NCBI GenBank SRA database (BioProject: PRJNA1468418; https://www.ncbi.nlm.nih.gov/bioproject/1468418; Accession numbers SAMN60281468-SAMN60281636) pending article acceptance. The R code, sample metadata, output files and data files used for analyses will be available in the Dryad Digital Repository pending article acceptance. <u>Benefit sharing</u>: A research collaboration was developed with scientists from Austria, Ecuador, Australia and the United States to collect samples and all collaborators are included as co-authors. The results of research have been shared with the international *Philornis downsi* Working Group, the Charles Darwin Research Station and Galapagos National Parks. This research addresses a priority concern to conserve land birds and control invasive species on the Galapagos Islands. Our group is committed to international scientific partnerships, as well as institutional capacity building.

## Additional files

Supplementary text, tables and figures are in the Supporting Information document.

## Author Contributions

This study was conceptualised by Sonia Kleindorfer, Rachael Y. Dudaniec, Lauren K. Common, Abbie C. Hay. Methodology was developed by Abbie C. Hay, Rachael Y. Dudaniec, Lauren K. Common, Sally Potter, Sonia Kleindorfer. Data collection was performed by Abbie C. Hay, Lauren K. Common, Rachael Y. Dudaniec, Sonia Kleindorfer, Jennifer A.H. Koop, George E. Heimpel, Sarah A. Knutie, Birgit Fessl, Lorraine Pérez-Beauchamp. Molecular laboratory work was conducted by Abbie. C. Hay and Jennifer A. H. Koop. Formal analysis designed and conducted by Abbie C. Hay, Rachael Dudaniec and Sally Potter. The manuscript was fully drafted by Abbie C. Hay with reviewing and editing by all authors.

