## Supplementary Material for "Asymmetric migration shapes genetic structure of the invasive avian vampire fly (*Philornis downsi*) across the Galápagos Islands"

**Supplementary Text**

**Text S1: Sampling methods**

Adult *P. downsi* were sampled from all Galápagos islands using McPhail traps (BioQuip Products, California, USA). McPhail traps were baited with 150 mL fermented papaya sugar mixture (as described in Common et al. 2022). A total of 80 traps were deployed per island, with 40 traps in the lowlands and 40 traps in the agricultural zone/highlands. Traps were placed at 20m intervals within the trapping area in each of the highland and lowland habitats and hung at 1.5-6m height from tree branches to encompass the flying height range of both male and female *P. downsi* (Kleindorfer et al. 2016). After 4-5 days, traps were collected with adult *P. downsi* flies identified in the field.

In addition, in March 2024, samples of *P. downsi* pupae were collected from terminated nests of the Little Vermilion Flycatcher *(Pyrocephalus nanus)* on Isabela Island following nest monitoring and parasite collection protocols outlined in Cimadom et al. (2014), and from nests of the Small Ground Finch (*Geospiza fuliginosa*) on San Cristóbal Island following the nest monitoring protocol outlined in Harvey et al. (2021) and parasite collection protocol outlined in Koop et al. (2013). Samples from mainland Ecuador were obtained from natural nests and artificial nest boxes hosted by four bird species (*Myiarchus phaeocephalus, Saltator striatipectus, Onychorhynchus coronatus, Troglodytes aedon musculus*), following the protocol of Bulgarella et al. (2015).

A total of 39 individuals across three sites on the Ecuadorian mainland were sampled. Mainland flies were reared to adulthood from *P. downsi* puparia collected from six nests at

Cerro Blanco (2–5 flies per nest; 3.9–12.5% of the total *P. downsi* intensity per nest), two nests at Agua Blanca (7 flies per nest; 17.9% and 25.0% of each nest) and one nest at Loma Alta (4 flies; 10.3% of the nest). A total of 235 individuals from across five Galápagos islands were sampled. This consisted of 138 individuals caught in traps (all individuals from Santa Cruz, Floreana and Santiago, 18.1% of individuals from Isabela, and 10.5% of individuals from San Cristóbal), and 97 individuals from nests (81.9% and 89.5% of individuals from Isabela and San Cristóbal respectively). For nest-reared samples from Isabela, between 1 and 4 adult flies were sampled per nest, reared from 21 nests (equivalent to 9.8-100% of the total *P. downsi* intensity per nest). For San Cristóbal, a total of five terminated nests were sampled for both adult flies and pupae, resulting in 1-12 flies sampled per nest (equivalent to 3.7-29.7% of the total *P. downsi* intensity per nest). We sampled a wide range of percentages of the total intensity across nests (3.7-100% overall) to both minimise the sampling of related individuals and to maximise capturing genetic diversity of unrelated individuals within nests. Dudaniec et al. (2010) found that at least 10% of the infra-population was sufficient to characterise genetic relatedness among the cohorts of *P. downsi* present within a nest, therefore all individuals collected from nests were analysed for genetic relatedness with full and half siblings removed prior to analysis (see Methods).

**Text S2: DNA extraction of *P. downsi***

DNA extractions were performed using DNeasy Blood & Tissue kits (Qiagen) using a modified manufacturers protocol. DNA was extracted from each adult fly using a single sample comprising the head, thorax, and 1–2 legs. The abdomen was excluded due to the presence of haploid sperm and eggs, which can complicate genetic sequencing results. For pupae, DNA was extracted from half of the individual (including tissue and pupal case).

Tissue samples from each *P. downsi* specimen were sliced into several pieces with a sterile scalpel in a sterile petri dish, placed in 1.5 mL microcentrifuge tubes with 180 µl lysis buffer ATL and lysed in a Qiagen Tissue Lyser for 3 mins at 40hz. Proteinase K (20µl) was added to each sample before vortexing for 10 seconds. Incubation was performed overnight at 56°C.

Lysis buffer AL (200µl) was added to each sample before vortexing for 10 seconds. Further incubation was performed at 56°C for 10 minutes. Proteins were precipitated with 200µl ethanol (100% absolute) and samples were vortexed for 10 seconds. Each sample was then briefly spun down in a minicentrifuge to bring all exoskeleton to the bottom of the tube. Using 200 µl pipette tips, the supernatant containing the DNA was gently transferred into a DNeasy Mini spin column and placed in a 2 mL collection tube and centrifuged at 8000 rpm for 1 min. The flow-through was discarded and the spin column was placed in a new 2 mL collection tube. DNA in spin columns was washed with 500µl wash buffer AW1, centrifuged at 8000 rpm for 1 min. The flow through was discarded and the spin column was placed in the same 2mL collection tube. DNA in spin columns was then washed with 500ul wash buffer AW2, centrifuged at 13000 rpm for 3 mins. The spin columns were transferred to new 1.5 mL microcentrifuge tubes. Elution was performed in two steps. 30µl of AE elution buffer was added to the centre of the spin column membrane, incubated for 1 minute at room temperature, and centrifuged for 1 minute at 8000 rpm. This step was repeated with a further 30µl of AE Buffer to result in 60µl purified DNA for each sample. DNA was quantified using a Qubit™ fluorometer with a dsDNA Qubit Assay Kit.

DNA samples were standardised to approximately 40 ng/µl and Illumina DNA Prep M libraries were prepared by the Australian Genome Research Facility (AGRF; Melbourne, Australia) following the manufacturer's protocol (Illumina, 2025). Individually indexed libraries were pooled and sequenced at AGRF using a low coverage whole genome 2x150 bp paired-end sequencing approach, with 65-96 samples per lane, across four lanes of an Illumina NovaSeq X Plus 10B (300 cycle).

**Text S3: Justification for use of BeXY**

BeXY is a Bayesian inference method that uses genotype likelihoods, making it well suited to low-coverage sequencing data. Notably, simulation studies have demonstrated that BeXY maintains high accuracy in individual karyotype assignment (>97%) even at very low read depths (~1000 total reads per individual; Caduff et al. 2024). Importantly, BeXY does not require prior knowledge of the species' sex determination system or the identity of sex-linked chromosomes, instead employing a continuous ploidy parameterization to account for mapping ambiguities (Caduff et al. 2024). These features make BeXY particularly appropriate for non-model organisms and datasets where reference-genome fragmentation and genotype uncertainty exists.

**Text S4: Validating BeXY scaffold classification.**

To validate BeXY scaffold classification, we calculated mean sequencing depth across autosomal and sex-linked scaffolds identified from BeXY and compared against the expected patterns for an XY system: autosomes are equal between males and females, males display half the coverage of females for X-linked scaffolds, and Y-linked scaffolds are absent in females (Palmer et al. 2019). We generated pooled Binary Alignment Map (BAM) files for sexes, which combine the mapped reads of multiple individuals into a single BAM, using SAMtools merge (Li et al. 2009). Pooled BAM files were generated for morphological males, morphological females, and also for XX/XY groups assigned in BeXY. For each pool, mean depth per scaffold was estimated using SAMtools depth, then normalized by the number of individuals. Normalized mean depth per scaffold was then averaged across inferred scaffold categories (autosome, X-linked, Y-linked) and visualised for groups using bar plots in ggplot2 v4.0.0 (Wickham, 2016) in R. Male-to-female and XY-to-XX depth ratios were also calculated and compared to the expected values for an XY system.

**Text S5: Calculating adjusted effective population size**

An equation by Waples et al. (2014) was utilised to calculate an adjusted $N_{b(Adj2)}$ from the raw $N_b$, correcting for overlapping generation times in *P. downsi* (Causton et al. 2019) by integrating two life-history parameters: adult lifespan (AL) and age at maturity for females (∝). The AL was calculated using an equation from Waples et al. (2014) which incorporates ∝ (6 days; Lahuatte et al. 2024), and maximum breeding age. As *P. downsi* females have sperm storage capabilities (Bulgarella et al. 2015) and gravid females are caught year-round, the maximum age of females wild caught by Bulgarella et al. (2022) was used as a proxy for maximum breeding age (324 days). The effective population size per generation ($N_{e(Adj2)}$) was derived from $N_{b(Adj2)}$, integrating the same two life-history parameters, following a second equation from Waples et al. (2014). Finally, a third formula was adapted from Waples et al. (2016) to control for physical linkage inherent in datasets with thousands of SNPs and few chromosomes, which can downwards bias $N_e$ estimates. We adjusted for the chromosome number for *Passeromyia heterochaeta,* 2n = 10 (Kaul and Tewari, 1979), using the haploid number, and present as $N_{e(Adj3)}$. As *Passeromyia* and *Philornis* form a monophyletic Muscid group (Couri and Carvalho, 2003), *P. heterochaeta* is, to the best of our knowledge, the closest karyotyped relative of *P. downsi.*

**Supplementary Tables**

**Table S1:** Filtering parameters and sites retained for the Galápagos only dataset. Filtering was performed using ANGSD (Korneliussen et al. 2014). Intermediate calculations were performed with other software: individual missing data and kinship via VCFtools (Danecek et al. 2011), autosomal scaffolds via BeXY (Caduff et al. 2024), linkage disequilibrium (LD) using ngsLD (Fox et al. 2019). Filtering thresholds were selected to balance marker retention and data quality e.g. Liu et al. (2024).

| Filtering step | N Ind | Average depth | ANGSD filtering thresholds | Additional info | Sites retained |
|---|---|---|---|---|---|
| Initial SNP call, to calculate missing data | 219 | 3.17x | -uniqueOnly 1 -remove_bads 1 -only_proper_pairs 1 -baq 1 -trim 0<br>-minMapQ 20 -minQ 20 -minInd 153 -setMinDepthInd 1 -setMaxDepth 2083 -doCounts 1<br>-GL 2 -doGlf 2 -doMajorMinor 1 -doMaf 1 -minMaf 0.02 -SNP_pval 1e-6 -rmTriallelic 0.05 \<br>-doPost 1 -doGeno 32 -doPlink 0 -doBcf 1 | minInd= 70% of total ind | 4951 |
| After exclusion of ind. with high missing data (>30%) | 189 | 3.31x | As above, except -minInd 95, -setMaxDepth 1877 | minInd= 50% of total ind | 70029 |
| Specifying autosomal scaffolds | 189 | 3.31x | As above, except -minInd 95, -setMaxDepth 1877, -rf autosomal scaffold file included | minInd= 50% of total ind | 66403 |
| **After exclusion of related ind. (>0.088)** | 169 | 3.27x | As above, except -minInd 85, -setMaxDepth 1658, -rf autosomal scaffold file included | minInd= 50% of total ind | 56063 |
| **HWE filtering (p > 0.001)** | 169 | 3.27x | As above, except -minInd 85, -setMaxDepth 1658, -rf autosomal scaffold file included, -minHWEpval 1e-6 | minInd= 50% of total ind | 55070 |
| **LD filtering ($r^2$ < 0.5 within 50 kb)** | 169 | 3.27x | Included -sites file to specify filtered and unlinked SNPs (dist 50kb, p-value 0.05) | minInd= 50% of total ind | 33638 |

**Table S2:** Filtering parameters and sites retained for the Galápagos and mainland Ecuador dataset. Filtering was performed using ANGSD (Korneliussen et al. 2014). Intermediate calculations were performed with other software: individual missing data and kinship via VCFtools (Danecek et al. 2011), autosomal scaffolds via BeXY (Caduff et al. 2024), linkage disequilibrium (LD) using ngsLD (Fox et al. 2019). Filtering thresholds were selected to balance marker retention and data quality e.g. Liu et al. (2024).

| **Filtering step** | **N Ind** | **Average depth** | **ANGSD filtering thresholds** | **Additional info** | **Sites retained** |
|---|---|---|---|---|---|
| Initial SNP call, to calculate missing data | 208 | 3.63x | -uniqueOnly 1 -remove_bads 1 -only_proper_pairs 1 -baq 1 -trim 0 -minMapQ 20 -minQ 20 -minInd 146 -setMinDepthInd 1 -setMaxDepth 2265 -doCounts 1 -GL 2 -doGlf 2 -doMajorMinor 1 -doMaf 1 -minMaf 0.02 -SNP_pval 1e-6 -rmTriallelic 0.05 -doPost 1 -doGeno 32 -doPlink 0 -doBcf 1 | minInd= 70% of total ind | 6667 |
| After exclusion of ind. with high missing data (>30%) | 198 | 3.66x | As above, except -minInd 99, -setMaxDepth 2174 | minInd= 50% of total ind | 179653 |
| Specifying autosomal scaffolds | 198 | 3.66x | As above, except -minInd 99, -setMaxDepth 2174, -rf autosomal scaffold file included | minInd= 50% of total ind | 169817 |
| **After exclusion of related ind. (>0.088)** | 185 | 3.57x | As above, except -minInd 93, -setMaxDepth 1981, -rf autosomal scaffold file included | minInd= 50% of total ind | 129130 |
| HWE filtering (p > **0.001)** | 185 | 3.57x | As above, except -minInd 93, -setMaxDepth 1981, -rf autosomal scaffold file included, -minHWEpval 1e-6 | minInd= 50% of total ind | 127883 |
| **LD filtering ($r^2$ < 0.5 within 50 kb)** | 185 | 3.57x | Included -sites file to specify filtered and unlinked SNPs (dist 50kb, p-value 0.05) | minInd= 50% of total ind | 77973 |

**Table S3:** Method of correction from effective number of breeders ($N_b$) to effective population size adjusted for overlapping generations and potential physical linkage bias ($N_{e(Adj3)}$), adapted from Bilgmann et al. (2021). Estimates of $N_{b(Adj2)}$, $N_{e(Adj2)}$ and $N_{e(Adj3)}$ were calculated using adjustments from Waples et al. (2014) and Waples et al. (2016). Adult life span (AL) for *P. downsi* was calculated using $\omega - \propto + 1$, where $\omega$ is the maximum breeding age (324 days; maximum age of females caught in Bulgarella et al. 2022) and $\propto$ is the age at first maturity for females (6 days; Lahuatte et al. 2024). Number of chromosomes (chr) for *Passeromyia heterochaeta* (2n = 10) (Kaul and Tewari 1978), close relative of *P. downsi* (Couri and Carvalho 2003), was incorporated using the haploid number.

| Formula | Source |
|---|---|
| $(1) N_{b(Adj2)} = \frac{N_b}{1.103 - 0.245 \times log\left(\frac{AL}{\propto}\right)}$ | Waples et al. (2014) |
| $(2) N_{e(Adj2)} = \frac{N_{b(Adj2)}}{0.485 + 0.758 \times log\left(\frac{AL}{\propto}\right)}$ | Waples et al. (2014) |
| $(3) N_{e(Adj3)} = \frac{N_{e(Adj2)}}{0.098 + 0.219 \times ln(chr)}$ | Waples et al. (2016) |

**Table S4:** Mean pairwise kinship of *P. downsi* within male and female groups on five Galápagos islands (Floreana, Isabela, San Cristóbal, Santa Cruz, Santiago). Values represent the mean kinship between all pairs of males and, separately, all pairs of females within each island. Inferred using the KING robust kinship estimator (Manichaikul et al. 2010) implemented in VCFtools relatedness2 (Danecek et al. 2011).

| Island | Females | Males |
|---|---|---|
| **Floreana** | 0.037 | 0.037 |
| **Isabela** | 0.040 | 0.041 |
| **San Cristóbal** | 0.055 | 0.057 |
| **Santa Cruz** | 0.042 | 0.042 |
| **Santiago** | 0.040 | 0.042 |

**Table S5:** Mean pairwise kinship of *P. downsi* males and females on five Galápagos islands to all individuals analysed on each island. For each sex, values represent the mean kinship of individual males/females to every other individual on the same island, averaged across all males or females. Inferred using the KING robust kinship estimator (Manichaikul et al. 2010) implemented in VCFtools relatedness2 (Danecek et al. 2011).

| Island | Females | Males |
|---|---|---|
| **Floreana** | 0.036 | 0.036 |
| **Isabela** | 0.039 | 0.039 |
| **San Cristóbal** | 0.051 | 0.052 |
| **Santa Cruz** | 0.041 | 0.040 |
| **Santiago** | 0.039 | 0.039 |

**Table S6:** Effective number of breeders ($N_b$) for *P. downsi* across the Galápagos archipelago and mainland Ecuador, estimated using NeEstimator2 (Do et al. 2014). $N_b$ was calculated across four critical values for rare alleles, using both parametric chi-squared 95% confidence intervals (CIs) and bias corrected JackKnife CIs (Waples & Do 2008). The values for Pcrit 0.05 are bolded and summarised in the main text. For individual Galápagos Islands (Isabela, San Cristóbal, Floreana, Santiago, Santa Cruz) a dataset of 33,609 SNPs was used. For Galápagos (including all five islands) and mainland Ecuador, a dataset of 30,142 SNPs was used. 'Four Island' refers to the genetic cluster of four islands (Isabela, Floreana, Santiago, San Cristóbal). Non-polymorphic loci per population were excluded, SNP count is reported.

| Location | Crit. Value | $N_b$ | Parametric CIs | | JackKnife CIs | |
|---|---|---|---|---|---|---|
| | | | L | U | L | U |
| Isabela (N = 54; 33,440 SNPs) | **0.05** | **139.2** | **139.1** | **139.4** | **114.1** | **176.6** |
| | 0.02 | -nan | Infinite | Infinite | Infinite | Infinite |
| | 0.01 | -nan | Infinite | Infinite | Infinite | Infinite |
| | +0 | 147.1 | 146.9 | 147.3 | 120.8 | 186.0 |
| San Cristóbal (N = 21; 29,025 SNPs) | **0.05** | **189.7** | **188.7** | **190.8** | **103.6** | **914.1** |
| | 0.02 | 215.2 | 214.0 | 216.4 | 113.2 | 1547.4 |
| | 0.01 | 215.2 | 214.0 | 216.4 | 113.2 | 1547.4 |
| | +0 | 215.2 | 214.0 | 216.4 | 113.2 | 1547.4 |
| Floreana (N = 27; 33,072 SNPs) | **0.05** | **199.0** | **198.3** | **199.8** | **147.4** | **302.4** |
| | 0.02 | 214.2 | 213.4 | 215.0 | 155.6 | 338.9 |
| | 0.01 | 224.6 | 223.7 | 225.5 | 161.4 | 363.6 |
| | +0 | 224.6 | 223.7 | 225.5 | 161.4 | 363.6 |
| Santiago (N = 33; 33,373 SNPs) | **0.05** | **213.6** | **213.0** | **214.2** | **165.7** | **297.5** |
| | 0.02 | 231.9 | 231.2 | 232.6 | 178.1 | 329.1 |
| | 0.01 | 239.4 | 238.7 | 240.2 | 182.7 | 343.7 |
| | +0 | 239.4 | 238.7 | 240.2 | 182.7 | 343.7 |
| Santa Cruz (N = 34; 33,367 SNPs) | **0.05** | **224.3** | **223.6** | **225.0** | **175.5** | **308.3** |
| | 0.02 | 240.1 | 239.4 | 240.9 | 185.7 | 336.8 |
| | 0.01 | 247.5 | 246.7 | 248.3 | 191.3 | 347.6 |
| | +0 | 247.5 | 246.7 | 248.3 | 191.3 | 347.6 |
| Four Island (N = 148; 33,587 SNPs) | **0.05** | **162.8** | **162.7** | **162.9** | **151.3** | **175.9** |
| | 0.02 | 169.0 | 168.9 | 169.1 | 157.1 | 182.4 |
| | 0.01 | 169.5 | 169.4 | 169.6 | 157.5 | 182.9 |
| | +0 | 169.6 | 169.5 | 169.7 | 157.7 | 183.1 |
| Galápagos (all) (N = 161; 22,940 SNPs) | **0.05** | **116.6** | **116.6** | **116.7** | **106.3** | **128.6** |
| | 0.02 | 123.7 | 123.6 | 123.7 | 112.7 | 136.4 |
| | 0.01 | 125.2 | 125.1 | 125.3 | 114.1 | 138.0 |
| | +0 | 134.2 | 134.1 | 134.2 | 122.4 | 147.7 |
| Mainland (N = 24; 27,883 SNPs) | **0.05** | **624.7** | **615.0** | **634.7** | **316.3** | **14095.2** |
| | 0.02 | 842.4 | 826.9 | 858.6 | 407.2 | Infinite |
| | 0.01 | 842.4 | 826.9 | 858.6 | 407.2 | Infinite |
| | +0 | 842.4 | 826.9 | 858.6 | 407.2 | Infinite |

**Table S7:** $N_b$ adjusted for overlapping generations ($N_{b\ (Adj2)}$), calculated using a formula in Waples et al. (2014) which incorporates maximum breeding age and female age at maturity. 'Four Island' refers to the genetic cluster of four islands (Isabela, Floreana, Santiago, San Cristóbal). The values for Pcrit 0.05 are bolded as these are summarised in the main text.

| Location | Crit. Value | $N_b$ (Adj2) | Parametric CIs | | JackKnife CIs | |
|---|---|---|---|---|---|---|
| | | | L | U | L | U |
| Isabela (N = 54; 33,440 SNPs) | **0.05** | **125.8** | **125.7** | **126.0** | **103.0** | **159.7** |
| | 0.02 | -nan | Infinite | Infinite | Infinite | Infinite |
| | 0.01 | -nan | Infinite | Infinite | Infinite | Infinite |
| | +0 | 132.9 | 132.8 | 133.1 | 109.1 | 168.2 |
| San Cristóbal (N = 21; 29,025 SNPs) | **0.05** | **171.6** | **170.7** | **172.6** | **93.5** | **828.3** |
| | 0.02 | 194.7 | 193.6 | 195.8 | 102.2 | 1402.5 |
| | 0.01 | 194.7 | 193.6 | 195.8 | 102.2 | 1402.5 |
| | +0 | 194.7 | 193.6 | 195.8 | 102.2 | 1402.5 |
| Floreana (N = 27; 33,072 SNPs) | **0.05** | **180.0** | **179.4** | **180.7** | **133.2** | **273.7** |
| | 0.02 | 193.8 | 193.0 | 194.5 | 140.6 | 306.8 |
| | 0.01 | 203.2 | 202.4 | 204.0 | 145.9 | 329.2 |
| | +0 | 203.2 | 202.4 | 204.0 | 145.9 | 329.2 |
| Santiago (N = 33; 33,373 SNPs) | **0.05** | **193.2** | **192.7** | **193.8** | **149.8** | **269.3** |
| | 0.02 | 209.8 | 209.2 | 210.5 | 161.0 | 297.9 |
| | 0.01 | 216.6 | 216.0 | 217.3 | 165.2 | 311.2 |
| | +0 | 216.6 | 216.0 | 217.3 | 165.2 | 311.2 |
| Santa Cruz (N = 34; 33,367 SNPs) | **0.05** | **202.9** | **202.3** | **203.6** | **158.7** | **279.1** |
| | 0.02 | 217.3 | 216.6 | 218.0 | 167.9 | 304.9 |
| | 0.01 | 224.0 | 223.2 | 224.7 | 173.0 | 314.7 |
| | +0 | 224.0 | 223.2 | 224.7 | 173.0 | 314.7 |
| Four Island (N = 148; 33,587 SNPs) | **0.05** | **147.2** | **147.1** | **147.3** | **136.7** | **159.1** |
| | 0.02 | 152.8 | 152.7 | 152.9 | 142.0 | 164.9 |
| | 0.01 | 153.2 | 153.2 | 153.3 | 142.4 | 165.4 |
| | +0 | 153.3 | 153.2 | 153.4 | 142.6 | 165.6 |
| Galápagos (all) (N = 161; 22,940 SNPs) | **0.05** | **105.3** | **105.3** | **105.4** | **96.0** | **116.2** |
| | 0.02 | 111.7 | 111.6 | 111.7 | 101.8 | 123.2 |
| | 0.01 | 113.1 | 113.0 | 113.2 | 103.0 | 124.7 |
| | +0 | 121.2 | 121.2 | 121.2 | 110.5 | 133.5 |
| Mainland (N = 24; 27,883 SNPs) | **0.05** | **565.9** | **557.1** | **575.0** | **286.3** | **12778.5** |
| | 0.02 | 763.3 | 749.3 | 778.0 | 368.8 | Infinite |
| | 0.01 | 763.3 | 749.3 | 778.0 | 368.8 | Infinite |
| | +0 | 763.3 | 749.3 | 778.0 | 368.8 | Infinite |

**Table S8:** Effective population size adjusted for overlapping generations ($N_{e(Adj2)}$), derived from $N_{b\ (Adj2)}$ using a formula from Waples et al. (2014) which incorporates maximum breeding age and female age at maturity. 'Four Island' refers to the genetic cluster of four islands (Isabela, Floreana, Santiago, San Cristóbal). The values for Pcrit 0.05 are bolded as these are summarised in the main text.

| Location | Crit. Value | $N_e$(Adj2) | Parametric CIs | | JackKnife CIs | |
|---|---|---|---|---|---|---|
| | | | L | U | L | U |
| Isabela (N = 54; 33,440 SNPs) | **0.05** | **260.6** | **260.5** | **261.0** | **213.7** | **330.6** |
| | 0.02 | -nan | Infinite | Infinite | Infinite | Infinite |
| | 0.01 | -nan | Infinite | Infinite | Infinite | Infinite |
| | 0 | 275.4 | 275.0 | 275.8 | 226.2 | 348.1 |
| San Cristóbal (N = 21; 29,025 SNPs) | **0.05** | **355.0** | **353.2** | **357.1** | **194.1** | **1709.2** |
| | 0.02 | 402.7 | 400.5 | 405.0 | 212.0 | 2893.0 |
| | 0.01 | 402.7 | 400.5 | 405.0 | 212.0 | 2893.0 |
| | 0 | 402.7 | 400.5 | 405.0 | 212.0 | 2893.0 |
| Floreana (N = 27; 33,072 SNPs) | **0.05** | **372.4** | **371.1** | **373.9** | **276.0** | **565.7** |
| | 0.02 | 400.8 | 399.3 | 402.3 | 291.3 | 633.9 |
| | 0.01 | 420.3 | 418.6 | 422.0 | 302.1 | 680.1 |
| | 0 | 420.3 | 418.6 | 422.0 | 302.1 | 680.1 |
| Santiago (N = 33; 33,373 SNPs) | **0.05** | **399.7** | **398.6** | **400.8** | **310.2** | **556.6** |
| | 0.02 | 433.9 | 432.6 | 435.2 | 333.4 | 615.6 |
| | 0.01 | 448.0 | 446.6 | 449.4 | 342.0 | 642.9 |
| | 0 | 448.0 | 446.6 | 449.4 | 342.0 | 642.9 |
| Santa Cruz (N = 34; 33,367 SNPs) | **0.05** | **419.7** | **418.4** | **421.0** | **328.5** | **576.7** |
| | 0.02 | 449.3 | 448.0 | 450.8 | 347.6 | 630.0 |
| | 0.01 | 463.1 | 461.6 | 464.6 | 358.0 | 650.2 |
| | 0 | 463.1 | 461.6 | 464.6 | 358.0 | 650.2 |
| Four Island (N = 148; 33,587 SNPs) | **0.05** | **304.8** | **304.6** | **304.9** | **283.3** | **329.2** |
| | 0.02 | 316.4 | 316.2 | 316.5 | 294.1 | 341.4 |
| | 0.01 | 317.3 | 317.1 | 317.5 | 294.9 | 342.3 |
| | 0 | 317.5 | 317.3 | 317.7 | 295.2 | 342.7 |
| Galápagos (all) (N = 161; 22,940 SNPs) | **0.05** | **218.4** | **218.4** | **218.6** | **199.1** | **240.8** |
| | 0.02 | 231.7 | 231.5 | 231.7 | 211.1 | 255.4 |
| | 0.01 | 234.5 | 234.3 | 234.7 | 213.7 | 258.4 |
| | 0 | 251.3 | 251.1 | 251.3 | 229.2 | 276.5 |
| Mainland (N = 24; 27,883 SNPs) | **0.05** | **1168.2** | **1150.1** | **1186.9** | **591.7** | **26348.8** |
| | 0.02 | 1575.1 | 1546.2 | 1605.4 | 761.6 | Infinite |
| | 0.01 | 1575.1 | 1546.2 | 1605.4 | 761.6 | Infinite |
| | 0 | 1575.1 | 1546.2 | 1605.4 | 761.6 | Infinite |

**Table S9:** Effective population size adjusted for physical linkage inherent in large SNP datasets $N_{e(Adj3)}$. Derived from $N_{e(Adj2)}$ using a formula rearranged from Waples et al. (2016), which incorporates chromosome number. 'Four Island' refers to the genetic cluster of four islands (Isabela, Floreana, Santiago, San Cristóbal). The values for Pcrit 0.05 are bolded as these are summarised in the main text.

| Location | Crit. Value | $N_e$(Adj3) | Parametric CIs | | JackKnife CIs | |
|---|---|---|---|---|---|---|
| | | | L | U | L | U |
| Isabela (N = 54; 33,440 SNPs) | **0.05** | **2660.0** | **2658.1** | **2663.8** | **2181.2** | **3373.4** |
| | 0.02 | -nan | Infinite | Infinite | Infinite | Infinite |
| | 0.01 | -nan | Infinite | Infinite | Infinite | Infinite |
| | 0 | 2810.7 | 2806.9 | 2814.5 | 2309.0 | 3552.7 |
| San Cristóbal (N = 21; 29,025 SNPs) | **0.05** | **3623.3** | **3604.2** | **3644.2** | **1980.9** | **17440.9** |
| | 0.02 | 4109.7 | 4086.8 | 4132.6 | 2164.1 | 29520.9 |
| | 0.01 | 4109.7 | 4086.8 | 4132.6 | 2164.1 | 29520.9 |
| | 0 | 4109.7 | 4086.8 | 4132.6 | 2164.1 | 29520.9 |
| Floreana (N = 27; 33,072 SNPs) | **0.05** | **3800.7** | **3787.3** | **3815.9** | **2816.4** | **5773.0** |
| | 0.02 | 4090.6 | 4075.3 | 4105.9 | 2972.8 | 6469.2 |
| | 0.01 | 4289.0 | 4271.8 | 4306.1 | 3083.5 | 6940.3 |
| | 0 | 4289.0 | 4271.8 | 4306.1 | 3083.5 | 6940.3 |
| Santiago (N = 33; 33,373 SNPs) | **0.05** | **4079.2** | **4067.7** | **4090.6** | **3165.5** | **5679.5** |
| | 0.02 | 4428.2 | 4414.9 | 4441.6 | 3402.0 | 6282.3 |
| | 0.01 | 4571.3 | 4557.9 | 4586.5 | 3489.7 | 6560.8 |
| | 0 | 4571.3 | 4557.9 | 4586.5 | 3489.7 | 6560.8 |
| Santa Cruz (N = 34; 33,367 SNPs) | **0.05** | **4283.2** | **4269.9** | **4296.6** | **3352.4** | **5885.5** |
| | 0.02 | 4584.6 | 4571.3 | 4599.9 | 3547.0 | 6429.1 |
| | 0.01 | 4725.8 | 4710.5 | 4741.0 | 3653.8 | 6635.2 |
| | 0 | 4725.8 | 4710.5 | 4741.0 | 3653.8 | 6635.2 |
| Four Island (N = 148; 33,587 SNPs) | **0.05** | **3110.2** | **3108.3** | **3112.1** | **2890.8** | **3360.0** |
| | 0.02 | 3228.4 | 3226.5 | 3230.3 | 3001.4 | 3484.0 |
| | 0.01 | 3238.0 | 3236.1 | 3239.9 | 3009.1 | 3493.6 |
| | 0 | 3239.9 | 3238.0 | 3241.8 | 3012.9 | 3497.4 |
| Galápagos (N = 161; 22,940 SNPs) | **0.05** | **2228.9** | **2228.9** | **2230.8** | **2032.4** | **2457.8** |
| | 0.02 | 2364.3 | 2362.4 | 2364.3 | 2154.5 | 2606.6 |
| | 0.01 | 2393.0 | 2391.0 | 2394.9 | 2181.2 | 2637.1 |
| | 0 | 2564.6 | 2562.7 | 2564.6 | 2339.5 | 2822.1 |
| Mainland (N = 24; 27,883 SNPs) | **0.05** | **11920.7** | **11735.7** | **12111.5** | **6038.1** | **268865.9** |
| | 0.02 | 16073.3 | 15777.6 | 16382.3 | 7772.0 | Infinite |
| | 0.01 | 16073.3 | 15777.6 | 16382.3 | 7772.0 | Infinite |
| | 0 | 16073.3 | 15777.6 | 16382.3 | 7772.0 | Infinite |

**Table S10:** Inferring sex-bias in putative migrants of *P. downsi* across five Galápagos islands (FL = Floreana, IS = Isabela, SO = Santiago, SZ = Santa Cruz, SL = San Cristóbal) based on individual assignments from WGSassign (DeSaix et al. 2024). Results are shown for three analyses: (a) unstandardized, (b) effective sample size (ESS) standardized by random subsampling, and (c) ESS standardized by subsampling individuals with the highest reference Z-score. Columns indicate the island of putative origin (highest assignment log-likelihood), and rows indicate the collection (reference) island. Cells report the number (proportion) of males (M) and females (F) assigned from the origin to the reference island as N_males (M_prop) / N_females (F_prop). Island codes not listed indicate no candidate migrants identified.

| ***a)*** | ***Unstandardized analysis*** | | | |
|---|---|---|---|---|
| | | *Reference island* | | |
| *Assigned island* | **FL (M / F)** | **IS (M / F)** | **SO (M / F)** | **SZ (M / F)** |
| **IS** | 3 (0.25) / 1 (0.07) | NA | 1 (0.08) / 2 (0.10) | 0 (0.00) / 2 (0.09) |
| **SO** | 1 (0.08) / 0 (0.00) | 1 (0.03) / 0 (0.00) | NA | 7 (0.64) / 7 (0.30) |
| **SZ** | 0 (0.00) / 0 (0.00) | 0 (0.00) / 1 (0.04) | 2 (0.17) / 6 (0.29) | NA |

| ***b)*** | ***Random standardization*** | | | |
|---|---|---|---|---|
| | | *Reference island* | | |
| *Assigned island* | **FL (M / F)** | **IS (M / F)** | **SO (M / F)** | **SZ (M / F)** |
| **FL** | NA | 3 (0.10) / 0 (0.00) | 1 (0.08) / 0 (0.00) | 0 (0.00) / 0 (0.00) |
| **IS** | 0 (0.00) / 0 (0.00) | NA | 0 (0.00) / 4 (0.19) | 0 (0.00) / 0 (0.00) |
| **SO** | 1 (0.08) / 0 (0.00) | 9 (0.31) / 1 (0.04) | NA | 5 (0.45) / 6 (0.26) |
| **SZ** | 0 (0.00) / 0 (0.00) | 0 (0.00) / 0 (0.00) | 3 (0.25) / 7 (0.33) | NA |

| **c)** | ***Standardization via reference Z-score*** | | | |
|---|---|---|---|---|
| | | *Reference island* | | |
| *Assigned island* | **FL (M / F)** | **IS (M / F)** | **SO (M / F)** | **SZ (M / F)** |
| **FL** | NA | 8 (0.28) / 0 (0.00) | 6 (0.50) / 2 (0.10) | 5 (0.45) / 0 (0.00) |
| **IS** | 1 (0.08) / 0 (0.00) | NA | 0 (0.00) / 1 (0.05) | 1 (0.09) / 0 (0.00) |
| **SO** | 0 (0.00) / 1 (0.07) | 1 (0.03) / 1 (0.04) | NA | 1 (0.09) / 9 (0.39) |
| **SZ** | 0 (0.00) / 0 (0.00) | 0 (0.00) / 1 (0.04) | 2 (0.17) / 6 (0.29) | NA |

**Table S11:** Relative genetic migration of *P. downsi* across five Galápagos islands, inferred using divMigrate (Sundqvist et al. 2016). Genetic distance was calculated using the *Nm* model (Alcala et al. 2014) using 1000 bootstrap replications. Significant asymmetric migration values are indicated by bold font and asterisks (*). Matrix displays relative migration in the direction from the island listed in column 1, to the island listed across row 1. All relative migration values are scaled such that the highest observed rate equals one, with remaining values representing proportions relative to this maximum; thus, the migration rates shown are not absolute, but reflect relative directional gene flow between islands.

| | **Isabela** | **San Cristóbal** | **Floreana** | **Santiago** | **Santa Cruz** |
|---:|---:|---:|---:|---:|---:|
| **Isabela** | NA | 0.14 | 0.63 | 0.85 | 0.74 |
| **San Cristóbal** | **0.30*** | NA | **0.27*** | **0.34*** | **0.32*** |
| **Floreana** | 0.70 | 0.16 | NA | 0.65 | 0.59 |
| **Santiago** | 0.88 | 0.16 | 0.61 | NA | 0.99 |
| **Santa Cruz** | 0.77 | 0.16 | 0.56 | 1.00 | NA |

**Supplementary Figures**

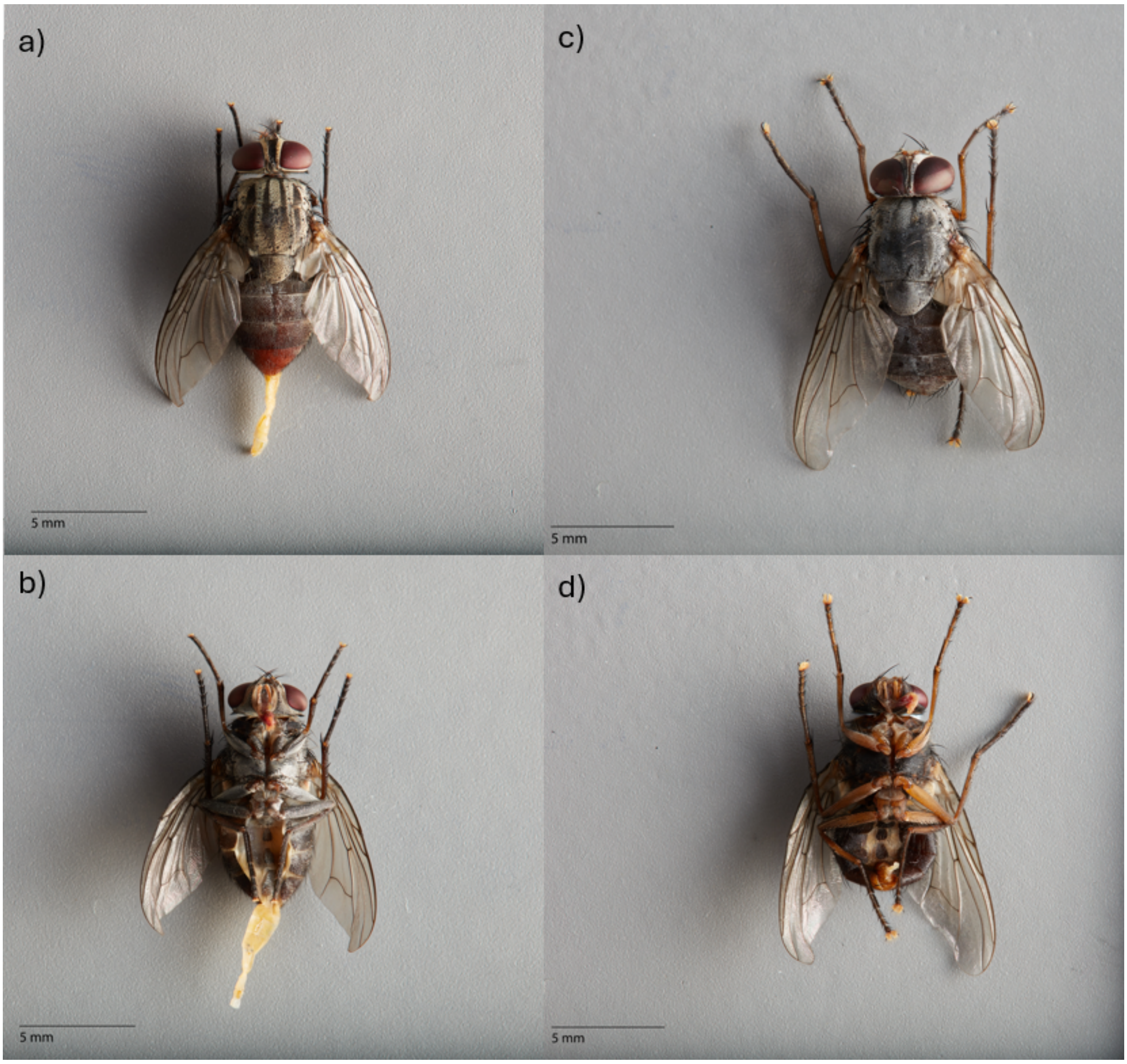


**Figure S1:** Examples of imaged *P. downsi* specimens collected across the Galápagos archipelago in 2024, a) female dorsal view, b) female ventral view, c) male dorsal view, d) male ventral view. All Galápagos specimens (n = 235) were imaged to verify sex identification. Legs of *P. downsi* are observed to be yellowish in males and blackish in females, with females exhibiting more widely set eyes and the ovipositor is visible following preservation in ethanol (Lahuatte et al. 2025; Fessl et al. 2001; Dodge and Aitken, 1968).

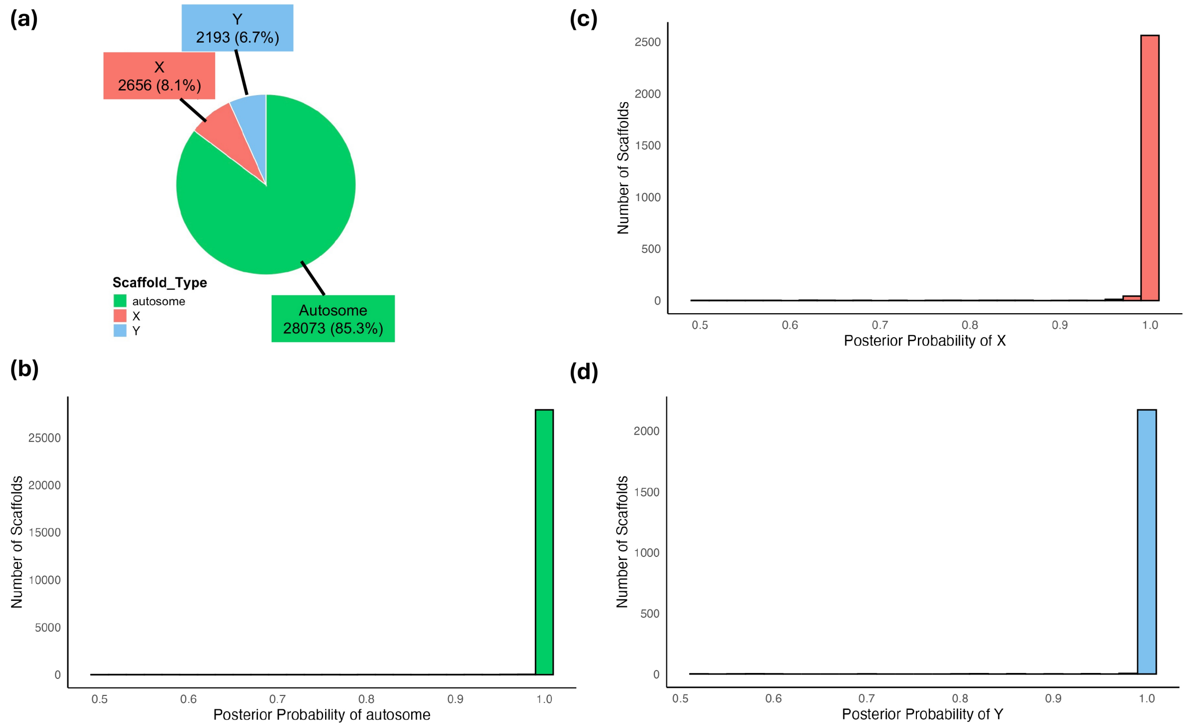


**Figure S2:** Assignment of scaffold types in *P. downsi* genome from 169 individuals collected across the Galápagos archipelago, inferred using BeXY (Caduff et al. 2024). Panel (a) depicts the number and percentages of all scaffolds analysed which are assigned with highest probability as X-linked, Y-linked and autosomal. Distribution of posterior probabilities for scaffolds assigned with highest confidence as (b) autosomal, (c) X-linked, and (d) Y-linked are presented.

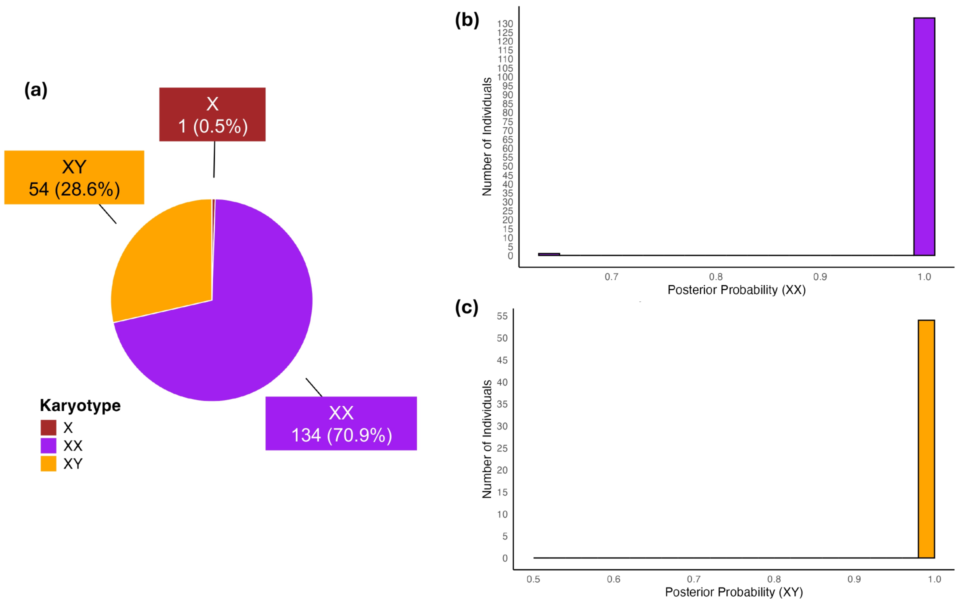


**Figure S3:** Assignment of individual sex karyotypes in *P. downsi* collected across the Galápagos, inferred using BeXY (Caduff et al. 2024). Panel (a) is the number and percentages of individuals assigned with the highest probability as genetic sex X, XX and XY. Distribution of posterior probability of assignment to karyotype with highest confidence for (b) XX inferred individuals, and (c) XY individuals are presented. One individual was assigned as the karyotype X with a posterior probability of 1.0.

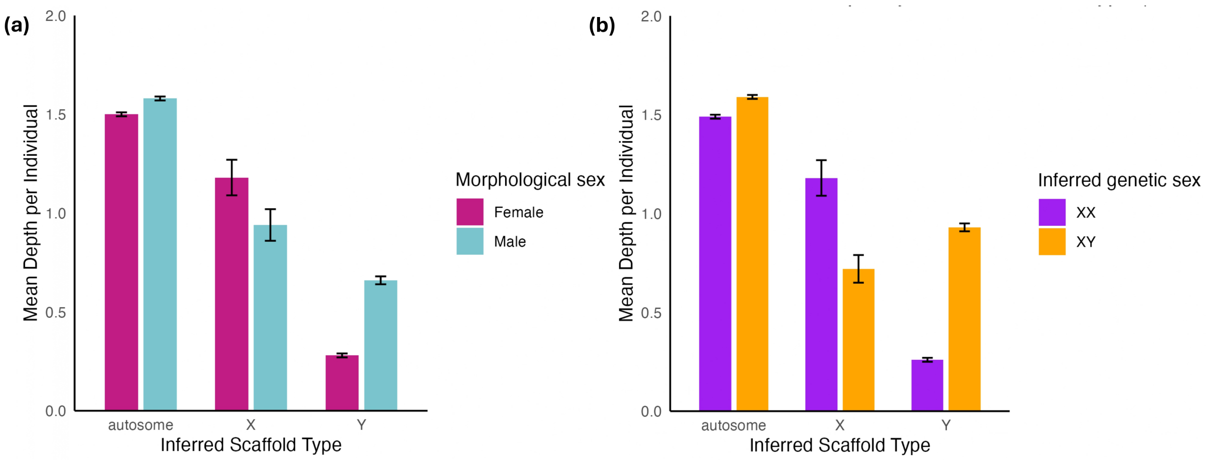


**Figure S4:** Mean depth of scaffold categories (autosome, X-linked, Y-linked) in *P. downsi* genome, inferred using BeXY (Caduff et al. 2024). Compared between (a) individuals identified as morphologically male and female, and (b) individuals genetically assigned as XX and XY in BeXY. Mean depth is normalized by the number of individuals in each group. Error bars represent mean ±1 standard error (SE) (SE = Standard deviation / $\sqrt{n}$, where n = number of scaffolds in the group).

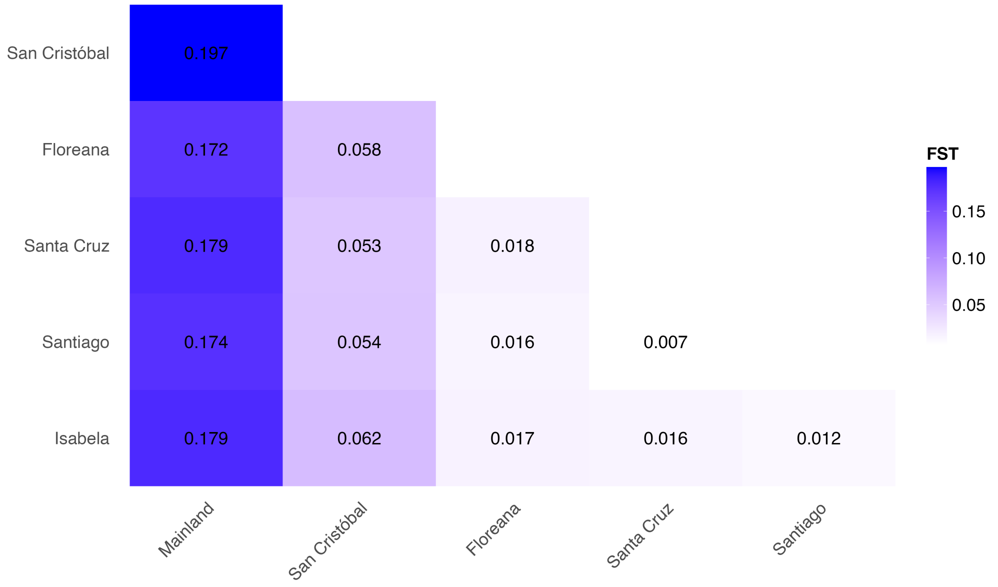


**Figure S5:** Global pairwise FST of *P. downsi* between each of the Galápagos islands and a reference group on mainland Ecuador (combined dataset). The mainland comparisons describe the combined global pairwise FST of individuals from across three sites on mainland Ecuador (Agua Blanca, Cerro Blanco and Loma Alta) with each Galápagos island.

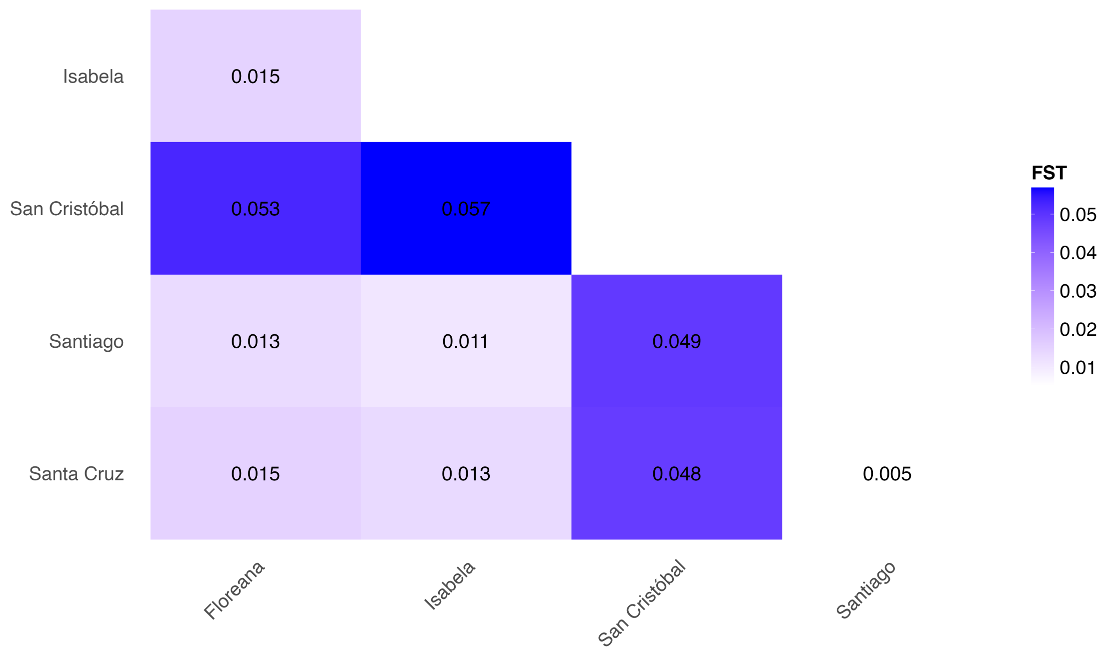


**Figure S6:** Global pairwise FST among *P. downsi* individuals collected across five Galápagos islands – Isabela, San Cristóbal, Santiago, Santa Cruz and Floreana. Computed using the Galápagos-only dataset (without mainland).

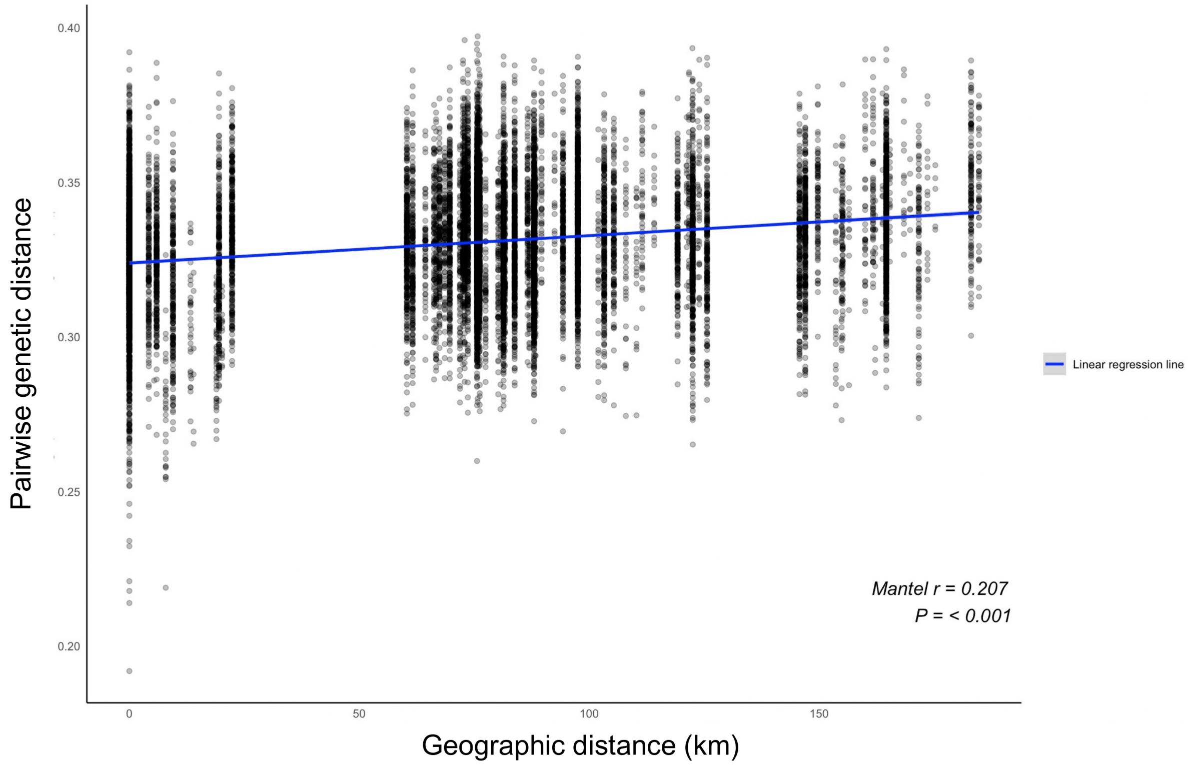


**Figure S7:** Isolation by genetic distance (pairwise geographic distance (km) vs genetic distance inferred using ngsDist; Vieira et al. 2016) of *P. downsi* individuals across five islands on the Galápagos archipelago (Galápagos-only dataset). Individuals are assigned to the geographic location of their sampling locality. Mantel r and p-value are indicated on the plot.

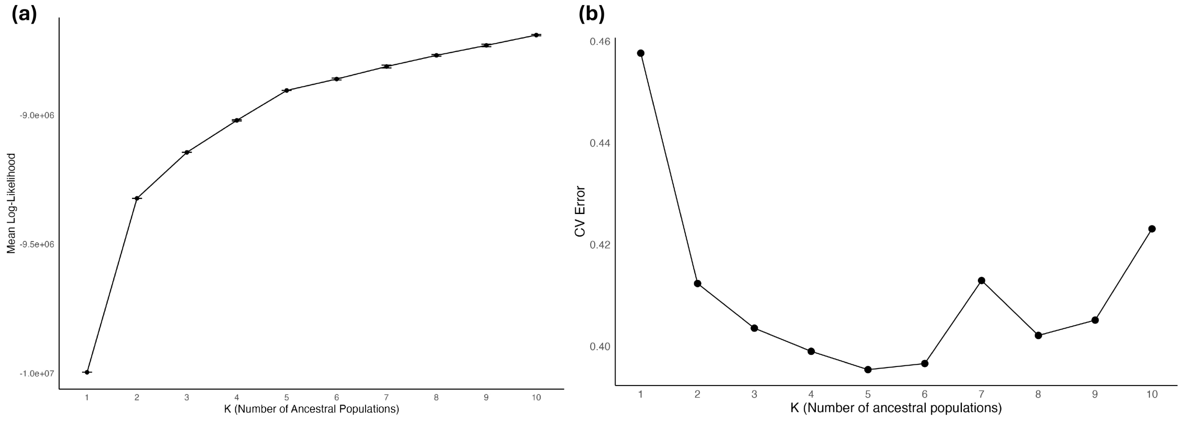


**Figure S8:** Comparison of most supported number of ancestral populations (K) for *P. downsi* across the Galápagos archipelago and mainland Ecuador, using genotype likelihood and called genotype methods, for K=1-10. Panel (a) depicts log-likelihoods computed using NGSadmix (Skotte et al. 2013) using 10 replicates of K, and (b) depicts cross-validation (CV) errors generated with ADMIXTURE (Alexander et al. 2009). Analysis is based on 77,973 SNPs.

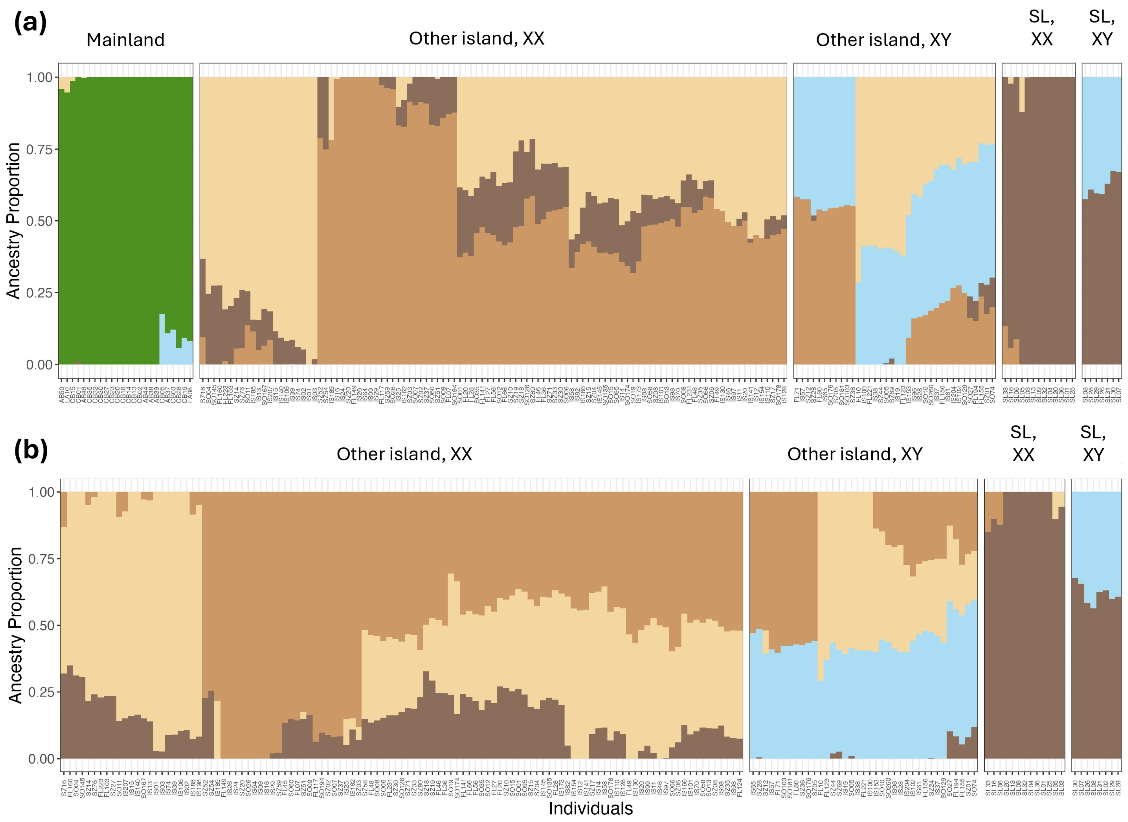


**Figure S9:** Admixture proportions for *P. downsi* individuals across (a) the Galápagos archipelago and mainland Ecuador using the combined dataset (K=5 best supported; 77,973 SNPs), and (b) the Galápagos archipelago using the Galápagos-only dataset (K=4 best supported; 33,638 SNPs). Admixture coefficients were calculated using ADMIXTURE (Alexander et al. 2009). The green cluster corresponds to ancestry from the mainland. Remaining admixture clusters do not associate with particular geographic locations, but individuals assigned as genetic sex XX and XY, across San Cristóbal Island (SL) and the other four islands (Floreana, Santa Cruz, Santiago, Isabela), do show distinct ancestry contributions.

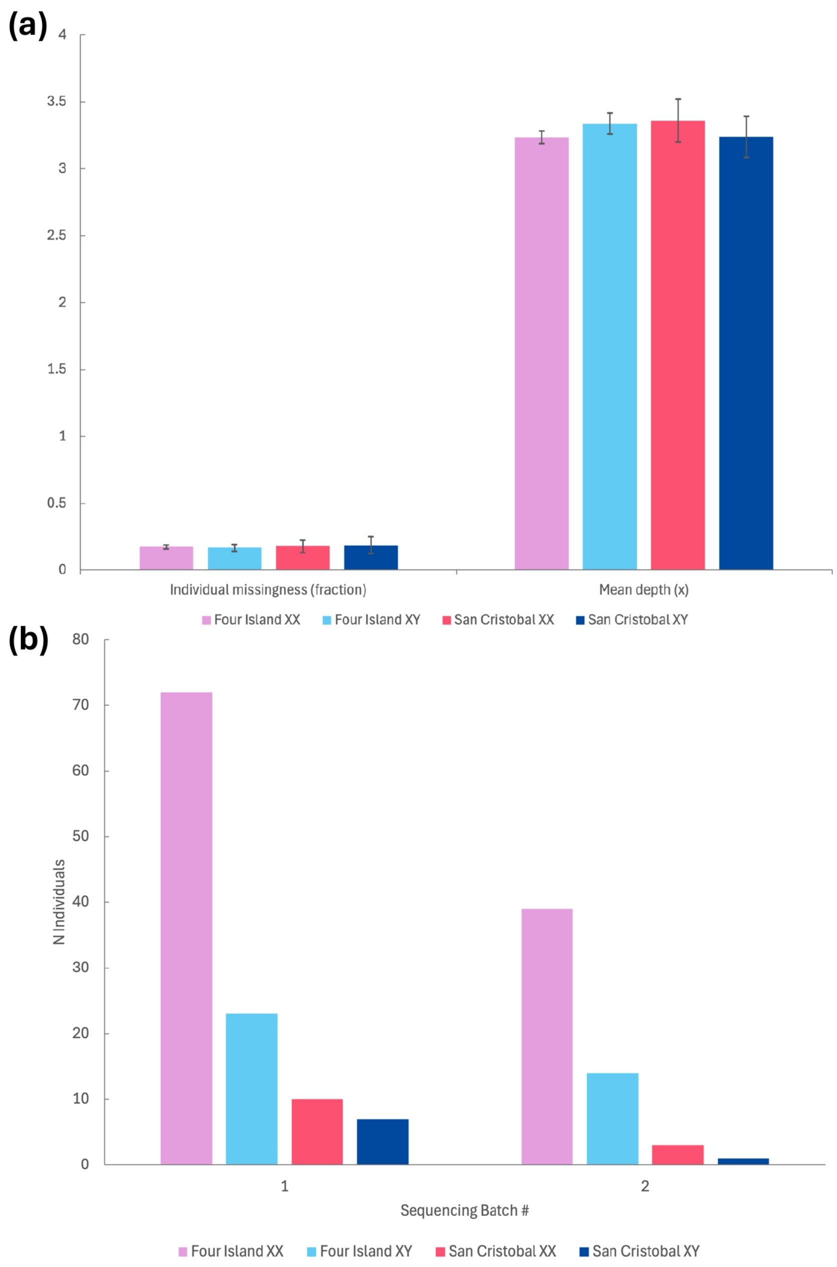


**Figure S10:** Comparison of potential technical and sequencing biases between PCA clusters inferred from the Galápagos-only dataset (Clusters: Four Island XX, Four Island XY, San Cristóbal XX, San Cristóbal XY). Panel a) depicts individual missingness (fraction of missing genotypes: 0-1) and mean estimated depth (x) compared between PCA clusters. Error bars represent standard error (SE). Panel b) depicts the count of individuals from separate Illumina sequencing runs (batch 1 or 2) appearing in PCA cluster groups. Batch 1 consists of 112 individuals and batch 2 consists of 57 individuals in total.

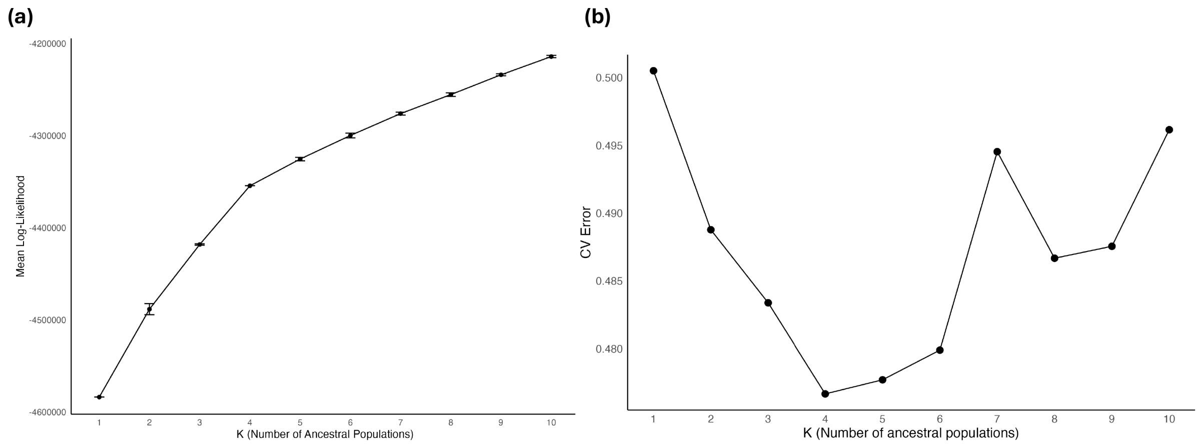


**Figure S11:** Comparison of most supported number of ancestral populations (K) for *P. downsi* across the Galápagos archipelago (Galápagos-only dataset), using genotype likelihood and called genotype methods, for K=1-10. Panel (a) depicts log-likelihoods computed using NGSadmix (Skotte et al. 2013) using 10 replicates of K, and (b) depicts cross-validation (CV) errors generated with ADMIXTURE (Alexander et al. 2009). Analysis is based on 33,638 SNPs.

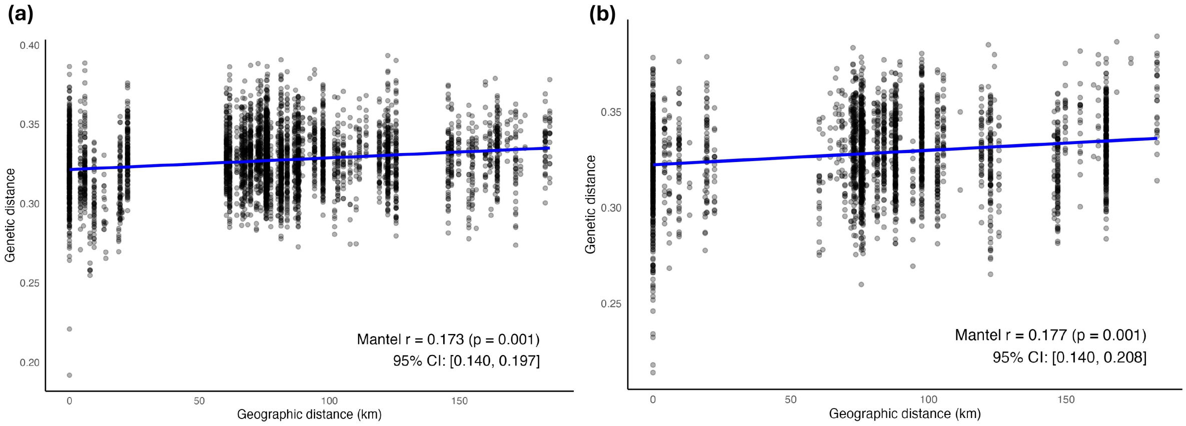


**Figure S12:** Isolation by genetic distance (pairwise geographic distance (km) vs genetic distance inferred using ngsDist; Vieira et al. 2016) of *P. downsi* individuals across five islands of the Galápagos archipelago (Galápagos-only dataset). Panel (a) analyses only morphologically identified females, and (b) analyses only morphologically identified males. Individuals are assigned to the geographic location of their sampling locality. Mantel r, p-value and 95% confidence interval (CI) are indicated on the plot.

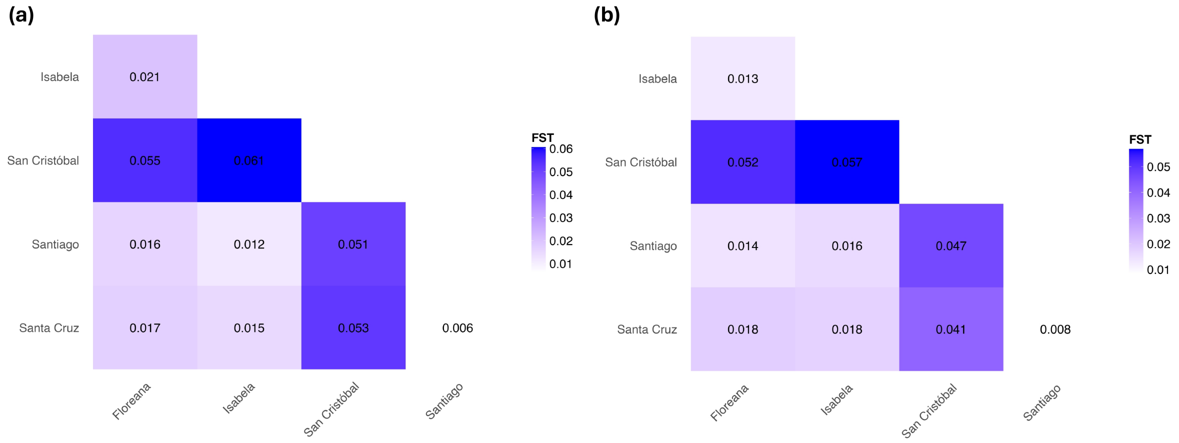


**Figure S13:** Global pairwise FST among *P. downsi* individuals collected across five Galápagos islands – Isabela, San Cristóbal, Santiago, Santa Cruz and Floreana (Galápagos-only dataset). Panel (a) compares morphologically identified females, and (b) compares morphologically identified males.

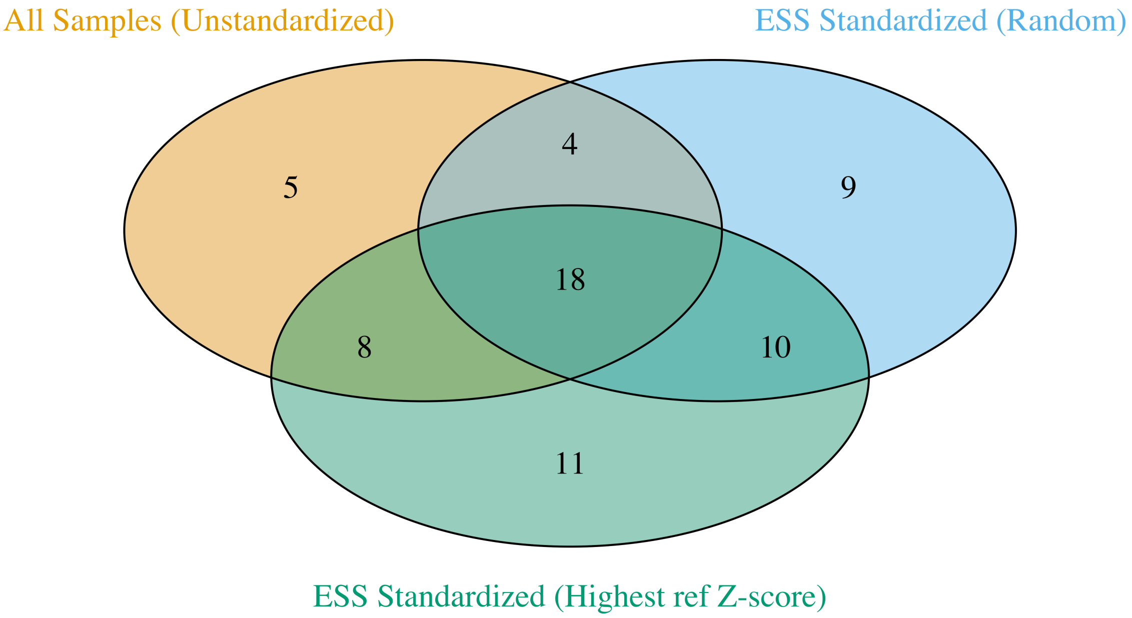


**Figure S14:** Comparison of candidate *P. downsi* migrants across the Galápagos Islands between three assignment approaches in WGSassign (DeSaix et al. 2024). Candidate migrants were identified as individuals that were assigned with highest likelihood to an island other than their sampled reference island. Three different methods were used to construct island reference populations in WGSassign; Left/yellow: all sampled individuals used (no effective sample size standardization); Right/blue: Effective sample size standardized via random subsampling per island; Lower/green: Effective sample size standardized via subsampling using highest reference Z-scores. The reference Z-score corresponds to assignment confidence for an individual to its reference island, as defined by WGSassign.
